# Land plant-specific H3K27 methyltransferases ATXR5 and ATXR6 control plant development and stress responses

**DOI:** 10.1101/2024.11.04.621985

**Authors:** Xiaoyi Li, Jie Pan, Huairen Zhang, Hui Li, Qian Liu, Danhua Jiang

## Abstract

The colonization of land by plants relies on numerous evolutionary innovations crucial for terrestrial adaptation. These include advances in sexual reproduction and the ability to properly respond to various environmental stresses, which involve precise control of their regulatory genes. A notable genetic innovation in land plants is the emergence of histone lysine methyltransferases ATXR5/6, which specifically catalyse the repressive histone H3 lysine 27 monomethylation (H3K27me1). However, the significance of the evolution of these enzymes in land plants remains unclear. In this study, we investigate the importance of ATXR5/6 by generating strong *atxr5;atxr6* double mutants in *Arabidopsis*. Our results show that ATXR5/6 are essential for plant reproductive development and play a critical role in supporting normal plant growth by repressing the transcription of stress responsive genes. In addition, ATXR5/6 are necessary for maintaining H3K27 trimethylation (H3K27me3), likely by providing H3K27me1 as a substrate for further methylation. We also demonstrate that the function of ATXR5/6 in regulating reproductive development and responsive genes is conserved in the monocot rice. We propose that land plants may have evolved ATXR5/6 to assist in the transcriptional regulation necessary for terrestrial adaptation.

## Introduction

The evolution of land plants from an ancestral charophycean alga establishes the path to modern terrestrial environment (Wodniok et al., 2011). During this process, plants have undergone numerous genetic innovations, such as the introduction of new gene families and the expansion of existing ones, to support their morphological and developmental changes, as well as the adaptation to environmental stresses (Bowles et al., 2020; Buschmann and Holzinger, 2020; Clark, 2023; Kumar et al., 2023). These developmental and environmental responsive genes often exhibit specific expression patterns rather than ubiquitous expression, suggesting that their expression needs to be tightly controlled to perform specialized functions in response to developmental and environmental signals (Aceituno et al., 2008).

In eukaryotes, epigenetic regulation, including post-translational modifications on histones, plays a pivotal role in determining the transcriptional activity (Zhao et al., 2019). Depending on their impact on transcription, histone modifications can either be “active” or “repressive”. A well-studied repressive histone modification is histone H3 lysine 9 (H3K9) methylation, which serves as a conserved marker of heterochromatin across yeast, animals, and plants (Montgomery et al., 2020; Naumann et al., 2005). In *Arabidopsis thaliana*, H3K9 methylation is primarily catalyzed by histone methyltransferases Su(var)3-9 homolog 4/Kryptonite (SUVH4/KYP), SUVH5 and SUVH6 (Du et al., 2015; Ebbs et al., 2005; Ebbs and Bender, 2006; Jackson et al., 2002), and it predominantly silences the transcription of transposable elements (TEs) enriched at heterochromatin regions (Ding et al., 2007; Ebbs *et al*., 2005; Ebbs and Bender, 2006; Vaillant and Paszkowski, 2007).

Another repressive histone modification, the histone H3 lysine 27 trimethylation (H3K27me3), is primarily enriched at euchromatin regions and is associated with inactive protein-coding genes (Zhang et al., 2007). In animals, the enzymes that catalyze H3K27me3 also deposit H3K27 monomethylation (H3K27me1) and H3K27 dimethylation (H3K27me2) (Ebert et al., 2004; Shen et al., 2008). However, in land plants, two plant-specific H3K27me1 methyltransferases, ARABIDOPSIS TRITHORAX-RELATED PROTEIN 5 (ATXR5) and ATXR6, have evolved (Supplemental Figure 1) (Jacob et al., 2009; Jacob and Michaels, 2009). H3K27me1 is mainly found at heterochromatin regions in *Arabidopsis* (Jacob *et al*., 2009; Mathieu et al., 2005). A weak *Arabidopsis atxr5;atxr6* double mutant exhibits decondensed heterochromatin, TE activation, and heterochromatin over-replication (Jacob *et al*., 2009; Jacob et al., 2010). Therefore, ATXR5/6-dependent H3K27me1 is regarded as a repressive histone modification that functions at heterochromatin in plants. However, it remains unclear why land plants evolved ATXR5/6 to deposit H3K27me1, given the existence of the already highly conserved heterochromatin marker, H3K9 methylation. Although some genes at euchromatin are found to carry H3K27me1 in *Arabidopsis* (Antunez-Sanchez et al., 2020; Potok et al., 2022), this genic H3K27me1 could predominantly result from the activity of H3K27 demethylases, such as REF6, ELF6 and JMJ13, which may demethylate H3K27me3 to produce H3K27me1 (Lu et al., 2011; Zheng et al., 2019). This hampers the efforts to identify the potential ATXR5/6-regulated genes through profiling H3K27me1 (Antunez-Sanchez *et al*., 2020; Potok *et al*., 2022). Thus, it is of great interest to directly determine the biological importance of ATXR5/6.

## Results

### Loss of ATXR5 and ATXR6 results in strong developmental defects

The so far commonly used *atxr5;atxr6* double mutant in *Arabidopsis* is an hypomorphic allele (referred to here as *atxr5;atxr6^hyp^*), in which *ATXR6* is still expressed at moderate levels because a T-DNA was inserted at its promoter region. The *atxr5;atxr6^hyp^* mutant displays only a slightly reduced growth phenotype compared with wild type (WT) Columbia (Col) (Jacob *et al*., 2009; Potok *et al*., 2022). To generate stronger mutants, we transformed CRISPR/Cas9 constructs targeting *ATXR6* into the *atxr5;atxr6^hyp^* mutant and created deletions within the genic region of *ATXR6* (*atxr6^c^-1* and *atxr6^c^-2*), which result in premature termination of translation (Figure 1A and Supplemental Figure 2A). We failed to recover any *atxr6^c^-1* or *atxr6^c^-2* homozygotes from the progeny of *atxr5;atxr5;atxr6^hyp^;atxr6^c^-1* or *atxr5;atxr5;atxr6^hyp^;atxr6^c^-2* (Supplemental Table 1), indicating that null *atxr5;atxr6* mutants are lethal. Examination of seeds in the siliques of *atxr5;atxr5;atxr6^hyp^;atxr6^c^-1* and *atxr5;atxr5;atxr6^hyp^;atxr6^c^-2* revealed seed abortion during seed development (Figure 1B). Moreover, small ovule-like structures were observed, which appeared to either not be fertilized or ceased development shortly after fertilization (Figure 1B). Reciprocal crosses indicated that the loss of *ATXR5* and *ATXR6* severely impaired female gametogenesis, although transmission was still observed at very low rates (Supplemental Table 2). Hence, we conclude that *atxr5;atxr6^c^-1* and *atxr5;atxr6^c^-2* mutants are not viable due to female gametophytic and embryonic lethality. This finding is consistent with recent reports showing that the *atxr5;atxr6* null mutants are lethal (Potok *et al*., 2022; Zhao et al., 2023).

**Figure 1.**
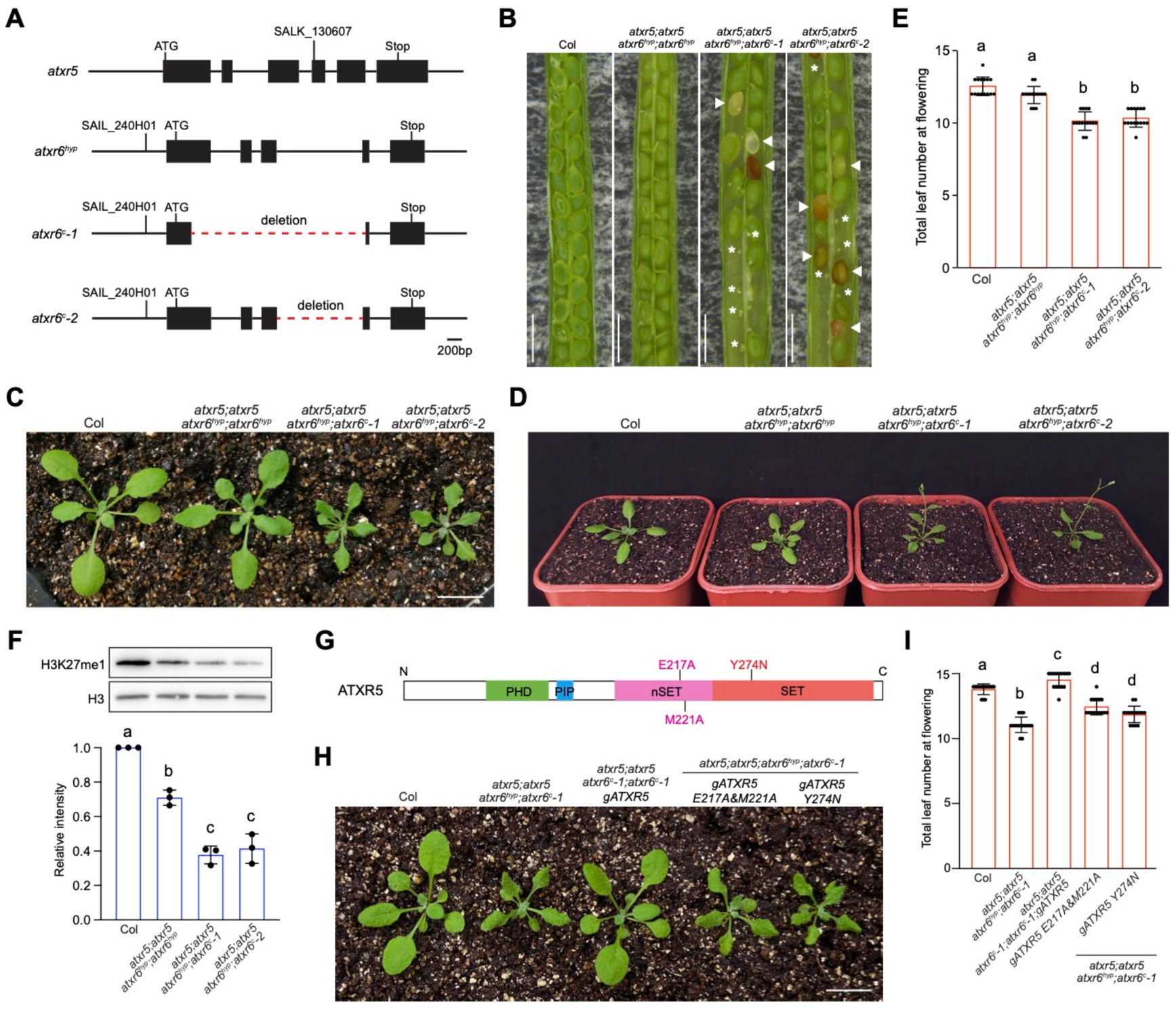
ATXR5 and ATXR6 regulate plant development. **A.** Schematic view of the full-length genomic structure of *ATXR5* and *ATXR6*. Filled boxes indicate exons, the red dashed lines represent deleted regions in *atxr6^c^-1* and *atxr6^c^-2*, SALK_130637 and SAIL_240H01 are T-DNA insertions. **B.** Seed developmental phenotypes of the indicated lines. Arrows indicate aborted seeds, while asterisks denote ovule-like structures. Scale bars, 1mm. **C.** Plant developmental phenotypes of the indicated lines. Scale bar, 1cm. **D.** Flowering phenotypes of the indicated lines. **E.** Total number of primary rosette and cauline leaves at flowering for the indicated lines. 15 plants were scored for each line. Values are means ± SD. The significance of differences was tested using one-way ANOVA with Tukey’s test (*P* < 0.05), with different letters indicating statistically significant differences. **F.** H3K27me1 levels in the indicated lines determined by western blotting. H3 was employed as a loading control. The bar chart represents the quantification of western blot signals from three biological replicates (Supplemental Figure 2C). Values are means ± SD. The significance of differences was tested using one-way ANOVA with Tukey’s test (*P* < 0.05), with different letters indicating statistically significant differences. **G.** Protein structure of ATXR5, the domains and point mutations generated to deactivate ATXR5 are presented. **H.** Plant developmental phenotypes of the complementation lines. Scale bar, 1cm. **I.** Total number of primary rosette and cauline leaves at flowering for the complementation lines. 15 plants were scored for each line. Values are means ± SD. The significance of differences was tested using one-way ANOVA with Tukey’s test (*P* < 0.05), with different letters indicating statistically significant differences.

Although the *atxr5;atxr6* null mutants are not viable, we observed that *atxr5;atxr5;atxr6^hyp^;atxr6^c^-1* and *atxr5;atxr5;atxr6^hyp^;atxr6^c^-2* displayed stronger developmental defects, such as significantly smaller plants, compared to *atxr5;atxr6^hyp^* at the vegetative stage (Figure 1C). Moreover, they exhibited early flowering phenotypes (Figure 1D and 1E). Examination of *ATXR6* expression revealed that its transcript levels were further reduced in *atxr5;atxr5;atxr6^hyp^;atxr6^c^-1* and *atxr5;atxr5;atxr6^hyp^;atxr6^c^-2* compared to *atxr5;atxr6^hyp^*(Supplemental Figure 2B), consistent with the more severe developmental defects observed in these mutants.

To determine whether ATXR5/6 regulate development via its enzymatic activity. We first analyzed global H3K27me1 levels in *atxr5;atxr5;atxr6^hyp^;atxr6^c^-1* and *atxr5;atxr5;atxr6^hyp^;atxr6^c^-2*. H3K27me1 levels were further diminished compared to *atxr5;atxr6^hyp^* (Figure 1F, and Supplemental Figure 2C), consistent with the more pronounced developmental defects. Next, we performed a complementation test with the full-length genomic *ATXR5* and the SET domain activity disrupted *ATXR5* (*gATXR5 E217A&M221A* and *gATXR5 Y274N*) (Figure 1G) (Jacob et al., 2014; Jacob *et al*., 2010). Only WT *ATXR5* fully rescued the developmental defects (Figure 1H and 1I and Supplemental Figure 2D), suggesting that the enzymatic activity of ATXR5/6 is crucial for plant development.

The function of ATXR5/6 in plant developmental control seems different from that of H3K9 methyltransferases, as loss of SUVH4, SUVH5, and SUVH6 does not induce significant developmental abnormalities (Supplemental Figure 3A) (Yu et al., 2017). The *atxr5;atxr6^hyp^* mutant bears excess DNA due to the over-replication of heterochromatin, particularly in endoreduplicated cells (Jacob *et al*., 2010). However, a comparable increase in DNA content was observed in *atxr5;atxr5;atxr6^hyp^;atxr6^c^-1* and *atxr5;atxr5;atxr6^hyp^;atxr6^c^-2* compared to *atxr5;atxr6^hyp^*(Supplemental Figure 3B and 3C), suggesting that the heterochromatin over-replication phenotype was not strongly enhanced in *atxr5;atxr5;atxr6^hyp^;atxr6^c^-1* and *atxr5;atxr5;atxr6^hyp^;atxr6^c^-2*. Moreover, similar percentages of nuclei in *atxr5;atxr6^hyp^*, *atxr5;atxr5;atxr6^hyp^;atxr6^c^-1* and *atxr5;atxr5;atxr6^hyp^;atxr6^c^-2* showed decondensation of chromocenters enriched with heterochromatin (Supplemental Figure 3D). These results suggest that the additional developmental defects observed in *atxr5;atxr5;atxr6^hyp^;atxr6^c^-1* and *atxr5;atxr5;atxr6^hyp^;atxr6^c^-2* compared to *atxr5;atxr6^hyp^*are likely not due to heterochromatin misregulation. Since *atxr5;atxr5;atxr6^hyp^;atxr6^c^-1* and *atxr5;atxr5;atxr6^hyp^;atxr6^c^-2* exhibited similar phenotypes, we mainly used *atxr5;atxr5;atxr6^hyp^;atxr6^c^-1* for subsequent studies.

### Loss of ATXR5 and ATXR6 enhances plant resistance to bacterial infection

To explore potential reasons for the additional developmental abnormalities in *atxr5;atxr5;atxr6^hyp^;atxr6^c^-1* compared to *atxr5;atxr6^hyp^*, we compared their transcriptome changes. Hundreds of TEs and genes were significantly misexpressed in *atxr5;atxr6^hyp^*and *atxr5;atxr5;atxr6^hyp^;atxr6^c^-1* compared to WT Col (fold change>2, *P*-adjust<0.05), with the majority showing increased transcript levels (Figure 2A) (Supplemental Dataset 1). This aligns with the notion that H3K27me1 is a repressive modification that inhibits transcription. *atxr5;atxr5;atxr6^hyp^;atxr6^c^-1* and *atxr5;atxr6^hyp^* had similar numbers of transcript level increased TEs, and nearly all of them were overlapped (Figure 2A and 2B and Supplemental Figure 4A). These results confirm our earlier observations that the *atxr5;atxr5;atxr6^hyp^;atxr6^c^-1* and *atxr5;atxr5;atxr6^hyp^;atxr6^c^-2* mutants did not show stronger defects at heterochromatin compared to *atxr5;atxr6^hyp^* (Supplemental Figure 3B-3D). On the other hand, many genes were significantly activated only in *atxr5;atxr5;atxr6^hyp^;atxr6^c^-1* but not in *atxr5;atxr6^hyp^* (Figure 2A and 2B and Supplemental Figure 4B). This suggests that the further loss of H3K27me1 in *atxr5;atxr5;atxr6^hyp^;atxr6^c^-1* mainly affects the repression of genes rather than TEs.

**Figure 2.**
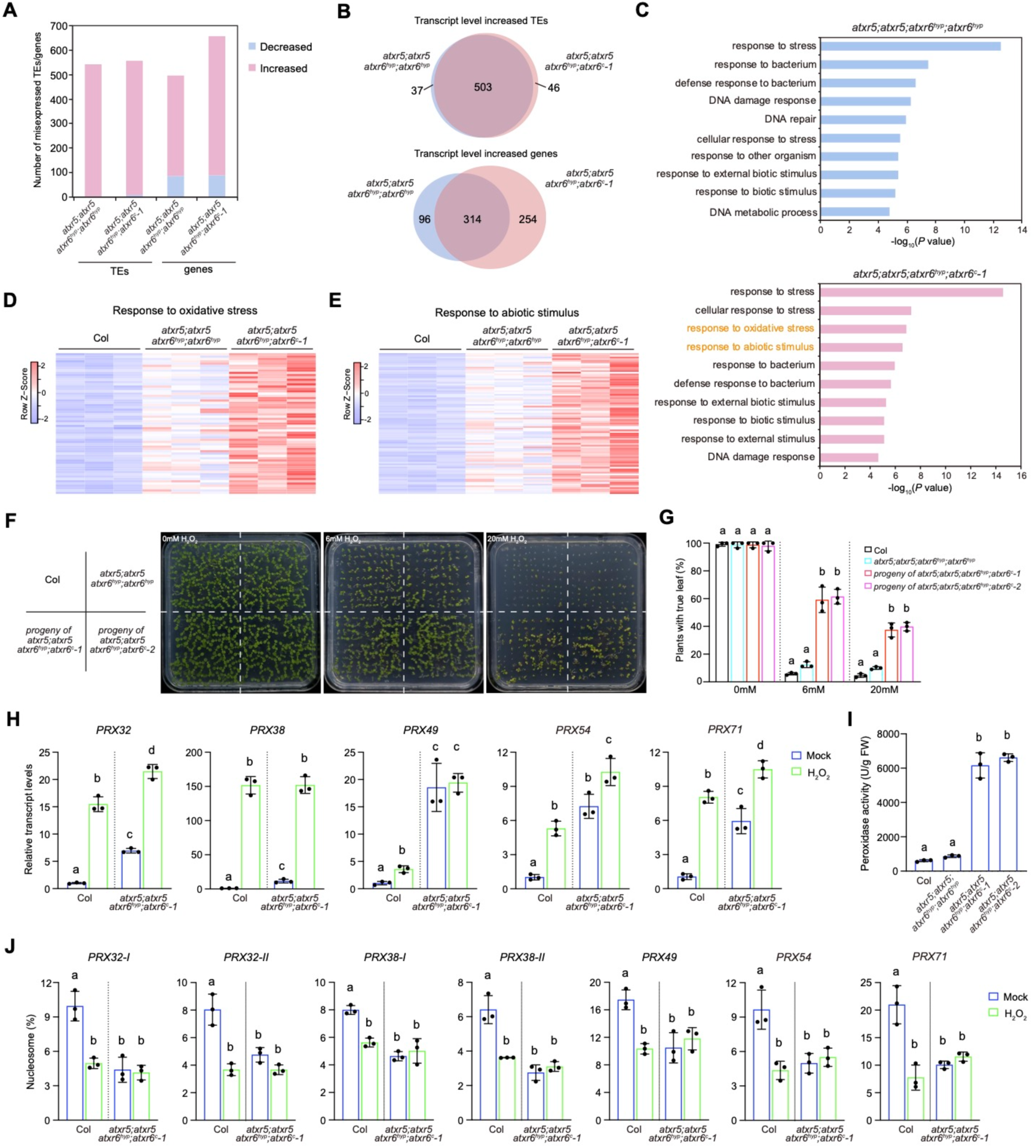
Loss of ATXR5 and ATXR6 induces stress responses. **A.** The number of significantly misexpressed TEs and genes in *atxr5;atxr6^hyp^* and *atxr5;atxr5;atxr6^hyp^;atxr6^c^-1*. **B.** Venn diagrams of transcript level significantly increased TEs and genes in *atxr5;atxr6^hyp^* and *atxr5;atxr5;atxr6^hyp^;atxr6^c^-1*. **C.** Gene ontology (GO) analysis on the transcript level significantly increased genes in *atxr5;atxr6^hyp^*and *atxr5;atxr5;atxr6^hyp^;atxr6^c^-1*. Top 10 representative terms are listed and ranked by *P* value. GO terms specifically enriched in *atxr5;atxr5;atxr6^hyp^;atxr6^c^-1* are highlighted in orange font. **D and E.** Heatmap showing transcript levels of oxidative stress response (d) and abiotic stimulus response (e) genes determined by RNA-seq. Results from three biological replicates are shown. **F.** Seedling growth phenotypes on 1/2MS plate supplemented with either no H_2_O_2_ or with 6mM or 20mM H_2_O_2_. Pictures were taken 10 days after germination. **G.** The true leaf formation rate of seedlings germinated on 1/2MS plate supplemented with either no H_2_O_2_ or with 6mM or 20mM H_2_O_2_. For the 0mM and 6mM H_2_O_2_ treatments, rates were measured 10 days after germination, while for the 20mM H_2_O_2_ treatment, rate were measured 15 days after germination. Values are means ± SD of three biological replicates. At least 79 seeds were sown for each replicate. The significance of differences was tested using one-way ANOVA with Tukey’s test (*P* < 0.05), with different letters indicating statistically significant differences. **H.** Transcript levels of peroxidase-coding genes determined by RT-qPCR in Col and *atxr5;atxr5;atxr6^hyp^;atxr6^c^-1* with or without H_2_O_2_ treatment. Values are means ± SD of three biological replicates. The significance of differences was tested using one-way ANOVA with Tukey’s test (*P* < 0.05), with different letters indicating statistically significant differences. **I.** Peroxidase activity in the indicated lines. Values are means ± SD of three biological replicates. The significance of differences was tested using one-way ANOVA with Tukey’s test (*P* < 0.05), with different letters indicating statistically significant differences. **J.** H3K27me1 levels at peroxidase-coding genes determined by ChIP-qPCR in Col and *atxr5;atxr5;atxr6^hyp^;atxr6^c^-1* with or without H_2_O_2_ treatment. The amounts of immunoprecipitated DNA fragments were normalized to input DNA (mononucleosome). Values are means ± SD of three biological replicates. The significance of differences was tested using one-way ANOVA with Tukey’s test (*P* < 0.05), with different letters indicating statistically significant differences.

We further performed gene ontology (GO) analysis with transcript level significantly increased genes in *atxr5;atxr6^hyp^* or *atxr5;atxr5;atxr6^hyp^;atxr6^c^-1* to assess the functions of ATXR5/6-repressed genes. Both groups were found to be enriched in pathways related to response to bacteria and defense response (Figure 2C and Supplemental Figure 4C). In addition, the DNA damage response pathway was also enriched (Figure 2C and Supplemental Figure 4D), consistent with previous studies showing that DNA damage repair genes are activated in *atxr5;atxr6^hyp^*, likely due to genome instability (Potok *et al*., 2022; Stroud et al., 2012). We next tested whether the loss of ATXR5/6-induced activation of defense responsive genes affects plant resistance to the virulent bacterium *P. syringae* pathovar tomato (*Pst*) DC3000. Compared to WT, less severe disease symptoms were observed on the leaves of *atxr5;atxr6^hyp^*, *atxr5;atxr5;atxr6^hyp^;atxr6^c^-1* and *atxr5;atxr5;atxr6^hyp^;atxr6^c^-2* three days after inoculation of *Pst* DC3000 (Supplemental Figure 4E). Consistently, bacterial growth was lower in these mutants compared to WT (Supplemental Figure 4F). However, it is important to note that the activation of DNA damage responses could enhance plant immunity (Wang et al., 2010; Yan et al., 2013). Moreover, both *atxr5;atxr6^hyp^*and *atxr5;atxr5;atxr6^hyp^;atxr6^c^-1* showed similar increase in DNA damage and defense responses, suggesting that these activations do not account for the additional developmental defects observed in *atxr5;atxr5;atxr6^hyp^;atxr6^c^-1* and *atxr5;atxr5;atxr6^hyp^;atxr6^c^-2*.

### ATXR5/6 repress plant oxidative stress response

Besides bacterial and DNA damage responses, GO analysis also revealed enrichment of oxidative stress and abiotic stimulus response pathways with transcript level significantly increased genes in *atxr5;atxr5;atxr6^hyp^;atxr6^c^-1* but not in *atxr5;atxr6^hyp^* (Figure 2C-2E), suggesting that the further loss of ATXR5/6 releases the repression of genes in these two pathways. To tested whether loss of ATXR5/6 affects plant resistance to oxidative stress, seeds collected from WT Col, *atxr5;atxr6^hyp^*, *atxr5;atxr5;atxr6^hyp^;atxr6^c^-1*, and *atxr5;atxr5;atxr6^hyp^;atxr6^c^-2* were germinated on 1/2MS plates containing hydrogen peroxide. Among the progeny of *atxr5;atxr5;atxr6^hyp^;atxr6^c^-1* and *atxr5;atxr5;atxr6^hyp^;atxr6^c^-2,* about half are expected to be *atxr5;atxr5;atxr6^hyp^;atxr6^c^-1* or *atxr5;atxr5;atxr6^hyp^;atxr6^c^-2*, respectively (Supplemental Table 1). By measuring the formation of true leaves, we found that the progeny of *atxr5;atxr5;atxr6^hyp^;atxr6^c^-1* and *atxr5;atxr5;atxr6^hyp^;atxr6^c^-2* exhibited greater resistance to hydrogen peroxide treatment compared to WT and the weak *atxr5;atxr6^hyp^* mutant (Figure 2F and 2G). Genotyping analysis confirmed that most plants forming true leaves under hydrogen peroxide treatment were indeed *atxr5;atxr5;atxr6^hyp^;atxr6^c^-1*or *atxr5;atxr5;atxr6^hyp^;atxr6^c^-2*. Similarly, *atxr5;atxr5;atxr6^hyp^;atxr6^c^-1* and *atxr5;atxr5;atxr6^hyp^;atxr6^c^-2* displayed increased resistance to salt stress (Supplemental Figure 5A and 5B), in agreement with the activation of abiotic stimulus response genes.

Notably, several class III peroxidase-coding genes were activated in *atxr5;atxr5;atxr6^hyp^;atxr6^c^-1* compared to WT (Figure 2D) (Supplemental Dataset 1). Overexpression of class III peroxidase-coding genes has been reported to cause reduced plant growth, a phenotype similar to that observed in both *atxr5;atxr5;atxr6^hyp^;atxr6^c^-1* and *atxr5;atxr5;atxr6^hyp^;atxr6^c^-2* (Pedreira et al., 2011; Raggi et al., 2015). In WT, the transcript levels of these peroxidase genes were induced by hydrogen peroxide treatment, suggesting that they are normally repressed in the absence of oxidative stress. However, their transcription was already activated in *atxr5;atxr5;atxr6^hyp^;atxr6^c^-1* even without stress (Figure 2H). Consistent with this, peroxidase activity was higher in *atxr5;atxr5;atxr6^hyp^;atxr6^c^-1* and *atxr5;atxr5;atxr6^hyp^;atxr6^c^-2* than in WT and *atxr5;atxr6^hyp^* (Figure 2I).

To examine whether ATXR5/6 repress the oxidative stress response through its enzymatic activity, we compared the oxidative stress response phenotypes of *atxr5;atxr5;atxr6^hyp^;atxr6^c^-1* complemented with either WT or SET domain mutated ATXR5. Only the WT ATXR5, and not the SET domain mutated ones, restored susceptibility to oxidative stress (Supplemental Figure 5C). Furthermore, enzymatic activity impaired ATXR5 was unable to repress the transcription of peroxidase-coding genes or reduce peroxidase activity (Supplemental Figure 5D and 5E). Thus, the repression of the oxidative stress response by ATXR5/6 is dependent on their enzymatic activity. We further analyzed H3K27me1 levels at these genes by ChIP-qPCR. H3K27me1 levels carried by these genes were reduced upon loss of ATXR5/6 (Figure 2J and Supplemental Figure 5F). Moreover, oxidative stress induced a decrease in H3K27me1 at these loci in WT (Figure 2J). Together, these results suggest that the ATXR5/6-mediated H3K27me1 represses peroxidase-coding genes under normal growth conditions.

### ATXR5/6 repress active histone modifications at peroxidase-coding genes

It was reported that ATXR5/6-mediated H3K27me1 antagonizes the deposition of active histone acetylation, such as H3K27 acetylation (H3K27ac) and H3K36ac, at the heterochromatin (Dong et al., 2021). We speculate that H3K27me1 may also inhibit other active modifications. At this point, we selected H3K27ac and H3K36 trimethylation (H3K36me3) to examine their levels at peroxidase-coding genes in *atxr5;atxr5;atxr6^hyp^;atxr6^c^-1*. Our results showed that H3K27ac and H3K36me3 levels were increased in *atxr5;atxr5;atxr6^hyp^;atxr6^c^-1* compared to WT (Figure 3A and 3B), suggesting that the ATXR5/6-mediated H3K27me1 represses their deposition at genes.

**Figure 3.**
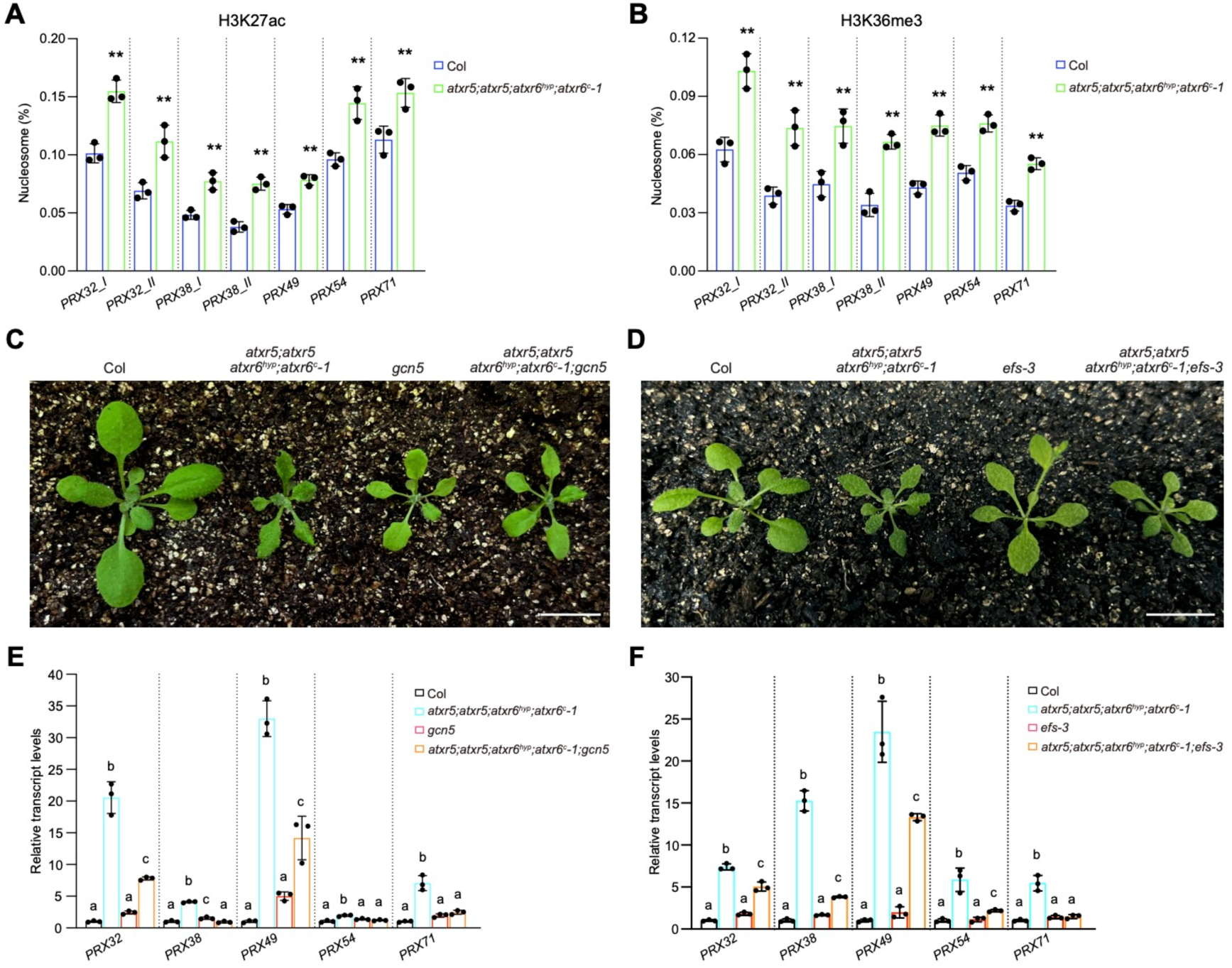
ATXR5 and ATXR6 antagonize the accumulation of H3K27ac and H3K36me3 at peroxidase-coding genes A and. **B.** H3K27ac (A) and H3K36me3 (B) levels at peroxidase-coding genes in Col and *atxr5;atxr5;atxr6^hyp^;atxr6^c^-1* determined by ChIP-qPCR. The amounts of immunoprecipitated DNA fragments were normalized to input DNA (mononucleosome). Values are means ± SD of three biological replicates. Statistical significance was determined by two-tailed Student’s *t*-test (**, *P* < 0.01). **C and D.** Plant developmental phenotypes following the loss of *GCN5* (C) or *EFS* (D) in *atxr5;atxr5;atxr6^hyp^;atxr6^c^-1*. Scale bars, 1cm. **E and F.** Transcript levels of peroxidase-coding genes following the loss of GCN5 (E) or EFS (F) in *atxr5;atxr5;atxr6^hyp^;atxr6^c^-1* determined by RT-qPCR. Values are means ± SD of three biological replicates. The significance of differences was tested using one-way ANOVA with Tukey’s test (*P* < 0.05), with different letters indicating statistically significant differences.

We then crossed *atxr5;atxr5;atxr6^hyp^;atxr6^c^-1* with a mutant of GENERAL CONTROL NON-REPRESSED PROTEIN 5 (GCN5) or EARLY FLOWERING IN SHORT DAYS (EFS, also known as SDG8), which are histone acetylase and methyltransferase responsible for H3K27ac and H3K36me3 deposition, respectively (Chen et al., 2017; Xu et al., 2008). The developmental phenotypes of the resulting triple mutants were closer to those of *gcn5* and *efs*, respectively (Figure 3C and 3D), suggesting that loss of GCN5 or EFS could partially repress the developmental defects induced by the ATXR5/6 mutation. Furthermore, the absence of GCN5 or EFS in *atxr5;atxr5;atxr6^hyp^;atxr6^c^-1* partially inhibited the activation of peroxidase-coding genes and reduced peroxidase activity (Figure 3E and 3F and Supplemental Figure 6). Therefore, the H3K27me1 deposited by ATXR5/6 likely prevents the accumulation of active histone modifications, thereby inhibiting gene activation.

### ATXR5/6 contribute to the maintenance of H3K27me3

The deposition of H3K27me1 by ATXR5/6 is considered replication coupled (Jacob *et al*., 2014; Raynaud et al., 2006). Similarly, the maintenance of another repressive histone mark, H3K27me3, is also closely linked with DNA replication (Del Olmo et al., 2016; Hyun et al., 2013; Jiang and Berger, 2017; Wang et al., 2023b; Zhou et al., 2017). It has been proposed that H3K27me1 deposition might facilitate the restoration of H3K27me3 after DNA replication (Borg et al., 2021; Jiang and Berger, 2017). To assess the requirement of ATXR5/6 in H3K27me3 maintenance, we performed ChIP-seq to examine H3K27me3 profiles in WT and *atxr5;atxr5;atxr6^hyp^;atxr6^c^-1*. Interestingly, we observed a moderate reduction in H3K27me3 levels at H3K27me3-enriched regions in *atxr5;atxr5;atxr6^hyp^;atxr6^c^-1* compared with WT (Figure 4A and 4B), and this reduction was consistently observed in both biological replicates performed (Supplemental Figure 7A). Similarly, H3K27me3 levels were lower at H3K27me3-enriched genes in *atxr5;atxr5;atxr6^hyp^;atxr6^c^-1* (Supplemental Figure 7B), and this was accompanied by a moderate increase in their transcription (Supplemental Figure 7C). The transcript levels of Polycomb group (PcG) genes, which regulate H3K27me3, as well as H3K27 demethylase-coding genes, were comparable between *atxr5;atxr5;atxr6^hyp^;atxr6^c^-1* and WT (Supplemental Figure 7D). Hence, the reduction of H3K27me3 in *atxr5;atxr5;atxr6^hyp^;atxr6^c^-1* is not due to the misexpression of these genes.

**Figure 4.**
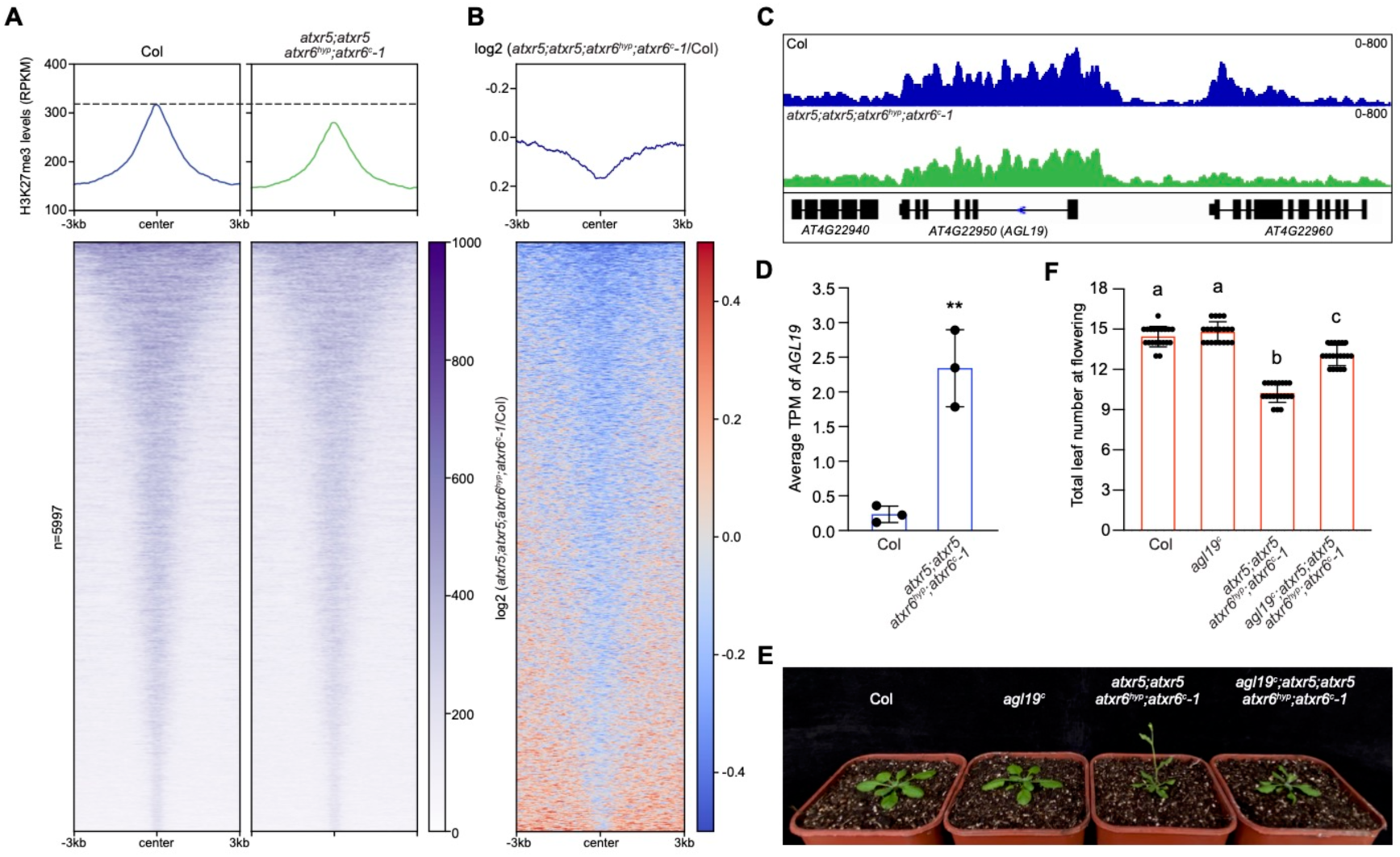
ATXR5 and ATXR6 are required for the maintenance of H3K27me3. **A.** Metaplots and heatmaps of H3K27me3 ChIP-seq signals in Col and *atxr5;atxr5;atxr6^hyp^;atxr6^c^-1* over H3K27me3-enriched peaks in WT Col. **B.** Changes in H3K27me3 ChIP-seq signals in *atxr5;atxr5;atxr6^hyp^;atxr6^c^-1* compared to Col over H3K27me3-enriched peaks in WT Col. **C.** Genome browser view of H3K27me3 accumulation levels at the locus encompassing *AGL19* in Col and *atxr5;atxr5;atxr6^hyp^;atxr6^c^-1*. **D.** Average TPM values of *AGL19* in Col and *atxr5;atxr5;atxr6^hyp^;atxr6^c^-1* determined by RNA-seq. Values are means ± SD of three biological replicates. Statistical significance was determined by two-tailed Student’s *t*-test (**, *P* < 0.01). **E.** Flowering phenotypes following loss of *AGL19* in *atxr5;atxr5;atxr6^hyp^;atxr6^c^-1*. **F.** Total number of primary rosette and cauline leaves at flowering for the indicated lines. 20 plants were scored for each line. Values are means ± SD. The significance of differences was tested using one-way ANOVA with Tukey’s test (*P* < 0.05), with different letters indicating statistically significant differences.

Both the *atxr5;atxr5;atxr6^hyp^;atxr6^c^-1* and *atxr5;atxr5;atxr6^hyp^;atxr6^c^-2* displayed early flowering phenotypes (Figure 1D and 1E). We noted that the floral activator *AGL19*, which is marked by H3K27me3, showed reduced H3K27me3 levels and increased transcript levels in *atxr5;atxr5;atxr6^hyp^;atxr6^c^-1* compared to WT (Figure 4C and 4D). In addition, the H3K27 monomethylation activity of ATXR5 is required for repressing both flowering and *AGL19* expression (Figure 1I and Supplemental Figure 8A). *AGL19* is strongly derepressed in PcG mutants, contributing to the early flowering phenotype in the loss of function mutant of CURLF LEAF (CLF), a key H3K27me3 methyltransferase in *Arabidopsis* (Schonrock et al., 2006). To determine whether the early flowering phenotype of *atxr5;atxr5;atxr6^hyp^;atxr6^c^-1* is due to upregulated *AGL19* expression, we generated an *agl19* mutant (*agl19^c^*) with CRISPR/Cas9 gene editing (Supplemental Figure 8B and 8C) and introduced it into the *atxr5;atxr5;atxr6^hyp^;atxr6^c^-1* background. Loss of *AGL19* partially suppressed the early flowering phenotype (Figure 4E and 4F), suggesting that ATXR5/6 repress flowering by assisting in the silencing of *AGL19*.

### OsATXR5/6 regulate reproductive development and repress responsive genes in rice

ATXR5 and ATXR6 are conserved proteins across land plants. We thus further investigated their function in the monocot model plant rice (*Oryza sativa*) by knocking out their coding genes with CRISPR/Cas9 (Supplemental Figure 9A-9D). Similar to what we found in *Arabidopsis*, we were unable to recover the *osatxr5^c^;osatxr6^c^*double homozygous plants from the progeny of *osatxr5^c^;osatxr6^c^/+* (Supplemental Table 3). While *osatxr5^c^;osatxr6^c^/+* produced normal flower organs and pollen (Supplemental Figure 9E and 9F), many empty grains were observed on the panicles, and its seed setting rate was lower than that of WT Nipponbare (NP) (Figure 5A and 5B). The ratio of *osatxr5^c^*to *osatxr5^c^;osatxr6^c^/+* in the progeny of *osatxr5^c^;osatxr6^c^/+* was approximately 1:2 (Supplemental Table 3). In addition, *osatxr5^c^;osatxr6^c^* mutations were successfully transmitted through both female and male gametes (Supplemental Table 4), suggesting that the *osatxr5^c^;osatxr6^c^* double homozygous is embryonic lethal. Thus, we conclude that OsATXR5/6 are essential for seed development but dispensable for gamete formation in rice.

**Figure 5.**
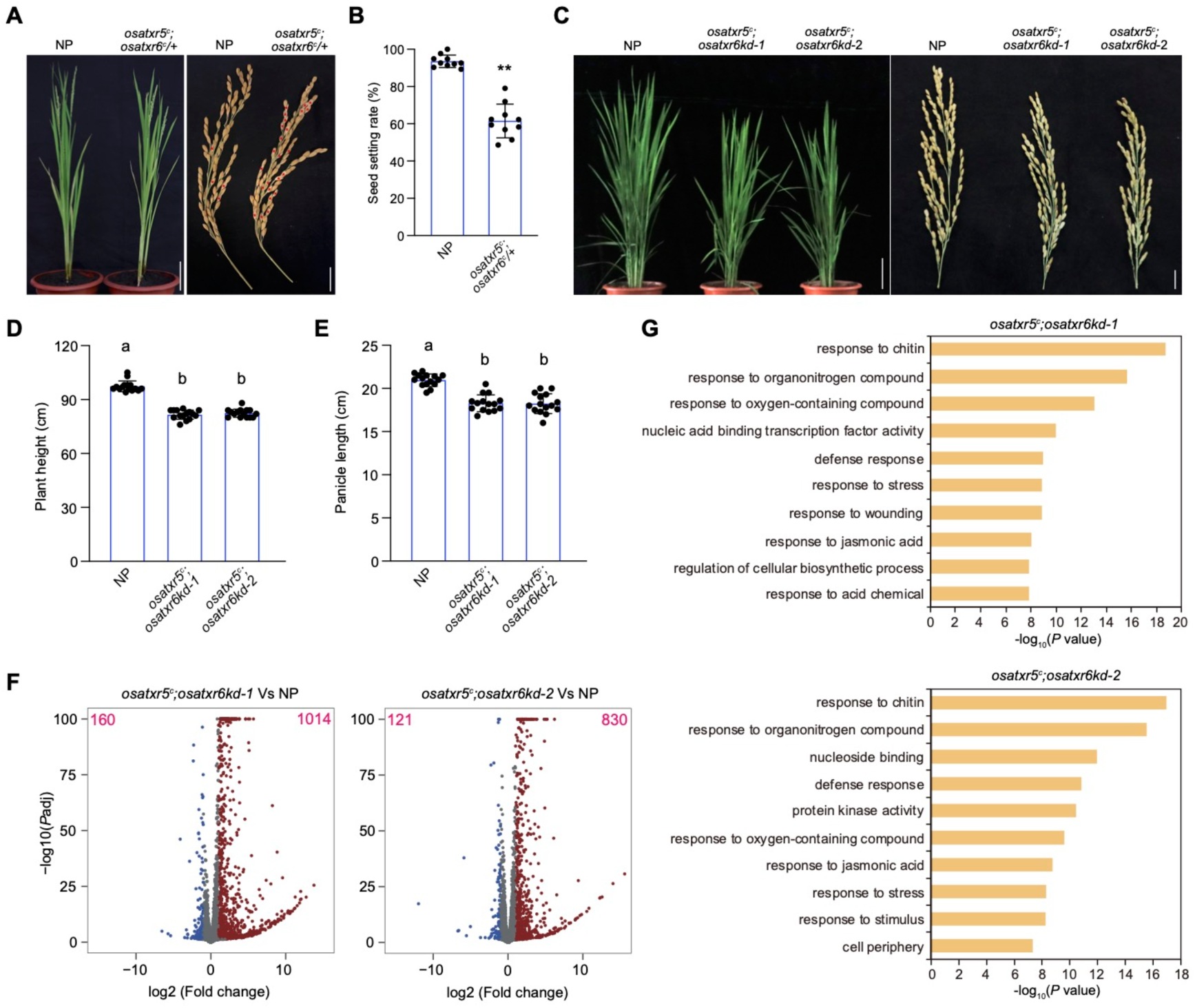
OsATXR5/6 are essential for reproductive development and the repression of responsive genes in rice. **A.** Plant (left) and panicle (right) phenotypes of NP and *osatxr5^c^;osatxr6^c^/+*. Empty grains are marked with red asterisks. Scale bars, 10cm (left) and 2cm (right). **B.** Seed set rates on the panicles of NP and *osatxr5^c^;osatxr6^c^/+*. 10 panicles were counted for each line. Values are means ± SD. Statistical significance was determined by two-tailed Student’s *t*-test (**, *P* < 0.01). **C.** Plant (left) and panicle (right) phenotypes of NP, *osatxr5^c^;osatxr6kd-1*, and *osatxr5^c^;osatxr6kd-2*. Scale bars, 10cm (left) and 2cm (right) **D and E.** Plant height (D) and panicle length (E) of NP, *osatxr5^c^;osatxr6kd-1*, and *osatxr5^c^;osatxr6kd-2*. 15 plants or panicles were scored for each line. Values are means ± SD. The significance of differences was tested using one-way ANOVA with Tukey’s test (*P* < 0.05), with different letters indicating statistically significant differences. **F.** Volcano plots of differentially expressed genes. The y-axis represents -log10 (*P* adjust), and the x-axis indicates log2 (fold change). Genes exhibiting at least a two-fold change in expression and an adjusted *P* value of less than 0.05 are considered misexpressed. The numbers of genes with increased and decreased transcript levels are indicated in the top right and left corners, respectively. **G.** Gene ontology (GO) analysis on transcript level significantly increased genes in *osatxr5^c^;osatxr6kd-1* and *osatxr5^c^;osatxr6kd-2*. Top 10 representative terms are listed and ranked by *P* value.

The *osatxr5^c^;osatxr6^c^/+* mutant did not show obvious developmental defects at the vegetative stage (Figure 5A). To assess the importance of OsATXR5/6 at this stage, we employed RNA interference (RNAi) strategy to knockdown *ATXR6* expression (*osatxr6kd*) in the *osatxr5^c^*mutant, and selected two independent lines for further studies (Supplemental Figure 10A). Both lines exhibited decreased global H3K27me1 levels and displayed reduced growth phenotypes compared to NP (Figure 5C-5E and Supplemental Figure 10B-10F). We next performed RNA-seq with NP, *osatxr5^c^;osatxr6kd-1*, and *osatxr5^c^;osatxr6kd-2*. Hundreds of TEs and genes were misexpressed in *osatxr5^c^;osatxr6kd-1* and *osatxr5^c^;osatxr6kd-2*, with the majority being upregulated (Figure 5F and Supplemental Figure 10G) (Supplemental Dataset 1), further supporting the role of OsATXR5/6 in transcriptional repression. GO analysis revealed that the derepressed genes were primarily enriched in responsive pathways (Figure 5G). Interestingly, DNA content analysis showed that the rice leaf predominantly contains 2C cells, and no extra DNA content was detected in *osatxr5^c^;osatxr6kd-1* and *osatxr5^c^;osatxr6kd-2* (Supplemental Figure 10H and 10I). Moreover, DNA damage response genes were not activated in *osatxr5^c^;osatxr6kd-1* and *osatxr5^c^;osatxr6kd-2* (Figure 5G) (Supplemental Dataset 1). Thus, unlike in *Arabidopsis*, the loss of OsATXR5/6 causes transcriptional activation without compromising DNA stability in rice. Together, these results suggest that ATXR5/6 have conserved roles in regulating development and responsive genes in both *Arabidopsis* and rice.

## Discussion

By generating strong *atxr5;atxr6* mutants in *Arabidopsis*, we have demonstrated that ATXR5/6 are crucial for regulating both plant development and abiotic stress responses. The function of ATXR5/6 in these processes relies on their enzymatic activity but is likely independent of their role in heterochromatin regulation, as the heterochromatin defects are comparable in the weak *atxr5;atxr6^hyp^* and the strong *atxr5;atxr5;atxr6^hyp^;atxr6^c^-1* and *atxr5;atxr5;atxr6^hyp^;atxr6^c^-2* mutants, with only the latter two showing clear developmental abnormalities and enhanced resistance to abiotic stresses (Figure 1 and 2, and Supplemental Figure 2-5). This aligns with the idea that heterochromatin defects in *Arabidopsis per se* do not typically cause strong developmental defects (Bourguet et al., 2021; Vongs et al., 1993). On the other hand, the further loss of ATXR5/6 in the strong *atxr5;atxr6* mutant induces the depression of many genes, including those responsive to oxidative stress and abiotic stimulus (Figure 2A-2E). ATXR5/6-mediated H3K27me1 is mainly enriched at heterochromatin regions (Antunez-Sanchez *et al*., 2020; Jacob *et al*., 2009; Potok *et al*., 2022). It is possible that a partial loss of ATXR5/6 in the weak *atxr5;atxr6^hyp^* mutant primarily impairs the deposition of H3K27me1 at heterochromatin, while a further loss of ATXR5/6 starts to cause defects at genic regions. This is further supported by the observations that complete loss of ATXR5/6 results in lethality, suggesting that a minimal level of ATXR5/6-mediated H3K27me1 is essential for plant viability.

Among the derepressed genes in the strong *atxr5;atxr6* mutant, there is a group of class III peroxidase-coding genes. Class III peroxidases are plant-specific peroxidases that can reduce peroxide and oxidize a wide range of substrates, such as lignin precursor (Passardi et al., 2004). Only a few class III peroxidases are found in some streptopyhte algae, but they have been greatly expanded in land plants (Mbadinga Mbadinga et al., 2020; Passardi *et al*., 2004). Therefore, class III peroxidases may play a critical role by adapting the land plants to a more oxygenated environment and enabling the formation of rigid cell walls suitable for terrestrial life (Mbadinga Mbadinga *et al*., 2020; Passardi et al., 2007; Passardi *et al*., 2004). As antioxidants, class III peroxidases are also important for plant responses to environmental stresses, with many being induced by stress signals (Eljebbawi et al., 2022; Kidwai et al., 2020). However, overexpression of class III peroxidases leads to reduced plant growth (Pedreira *et al*., 2011; Raggi *et al*., 2015), suggesting that their expression must be tightly controlled. It is likely that ATXR5/6 repress class III peroxidase-coding genes under normal growth conditions to balance plant growth and stress response.

We found that a loss of H3K27me1 at class III peroxidase-coding genes is accompanied by an increase in H3K27ac and H3K36me3 (Figure 2J, Figure 3A and 3B), suggesting that H3K27me1 represses active histone modifications. Introducing GCN5 and EFS mutations into the *atxr5;atxr5;atxr6^hyp^;atxr6^c^-1* mutant partially suppressed the activation of class III peroxidase-coding genes (Figure 3E and 3F). These observations are consistent with a recent study that reported an increase in H3K27ac at heterochromatin leading to heterochromatin defects and TE activation in *atxr5;atxr6^hyp^* (Dong *et al*., 2021). Therefore, AXTR5/6-mediated H3K27me1 appears to play a similar role in maintaining transcriptional silencing at both heterochromatin and genic regions by preventing the accumulation of active histone modifications.

In addition to depositing H3K27me1, we show that ATXR5/6 are also required for the maintenance of H3K27me3, a crucial histone modification in determining cellular identify (Figure 4 and Supplemental Figure 7) (Wiles and Selker, 2017). This observation supports the previous hypothesis that the ATXR5/6-deposited H3K27me1 during DNA replication may serve as a substrate for the further catalyzation and rapid recovery of H3K27me3 after DNA replication in plants (Borg *et al*., 2021; Jiang and Berger, 2017). Notably, H3K27me3 is only mildly affected in the strong *atxr5;atxr6* mutant. This may due to the residual ATXR6 still providing some H3K27me1, or the H3K27me3 methyltransferases may be capable of synthesizing most H3K27me3 directly from unmethylated H3K27. Nevertheless, this mild loss of H3K27me3 leads to moderate depression of H3K27me3-marked genes (Supplemental Figure 7C), and, in particular, the flowering time regulatory gene *AGL19* becomes strongly activated upon loss of ATXR5/6, which contributes to the accelerated floral transition phenotype of the strong *atxr5;atxr6* mutant, suggesting the importance of the ATXR5/6-assisted H3K27me3 maintenance in controlling development and cellular identity. We propose that land plants may have evolved ATXR5/6 for the strict maintenance of H3K27me3, which is necessary to support their complex developmental programs and the great diversity of their cell types. Additional studies are needed to distinguish the effects of ATXR5/6-dependent H3K27me1 and H3K27me3 in plant development and stress responses.

To test the function of ATXR5/6 in other plants, we generated the null *osatxr5;osatxr6* mutant in rice and found that OsATXR5/6 are also essential for plant reproduction (Figure 5A and 5B). However, unlike in *Arabidopsis*, OsATXR5/6 are required only for seed development and are dispensable for female gametogenesis. Further studies are needed to dissect the detailed functions of ATXR5/6 in plant reproductive development. In addition, by knocking down the expression of *OsATXR5/6*, we observed that the loss of OsATXR5/6 also leads to reduced plant growth and the depression of TEs and responsive genes (Figure 5C-5G and Supplemental Figure 10C-10G), suggesting the conserved function of OsATXR5/6 in repressing transcription and balancing plant growth and stress responses in rice. In *Arabidopsis*, the loss of ATXR5/6 results in genome instability, characterized by excess accumulation of heterochromatic DNA in endoreduplicated cells (Jacob *et al*., 2010). Interestingly, rice leaf cells lack endoreduplication, and no extra DNA was detected in the leaf nuclei of the *osatxr5;osatxr6* knockdown mutants (Supplemental Figure 10H and 10I). This suggests that loss of ATXR5/6 induces transcriptional activation independent from affecting genome instability, at least in rice.

In conclusion, this study demonstrates that ATXR5/6 play conserved roles in controlling plant development and stress responses, which may be essential for terrestrial adaptation. This provides a possible explanation for the emergence of ATXR5/6 in land plants. Furthermore, our work highlights complex functions of ATXR5/6 in regulating transcriptional repression at both heterochromatin and genic regions through mediating H3K27me1 and H3K27me3. Further studies are required to clarify these functions in more detail.

## Materials and Methods

### Plant materials and growth conditions

The *Arabidopsis atxr5;atxr6^hyp^* (Jacob *et al*., 2009)*, gcn5* (Salk_150784) (Kornet and Scheres, 2009), and *efs-3* (Kim et al., 2005) have been previously described. *Arabidopsis* plants were grown under long day conditions (16h light/8h dark) at 22°C. Rice plants were grown in paddy fields in Beijing, China.

### Plasmid construction for plant transformation

CRISPR-Cas9 gene editing constructs containing two single-guide RNAs targeting *Arabidopsis ATXR6* or *AGL19* were generated using pHSE401 as previously described (Xing et al., 2014). Constructs for editing *OsATXR5* and *OsATXR6* were generated using pYLCRISPR/Cas9P_ubi_-H, following the methods described by (Ma et al., 2015). Guide RNA sequences are listed in Supplemental Table 5. All the mutations were confirmed via Sanger sequencing. For the complementation test, the genomic DNA of *ATXR5*, including its promoter, was inserted into the pHGW vector (Karimi et al., 2002). The *ATXR5* sequence was further mutated to generate *gATXR5 E217A&M221A* or *gATXR5 Y274N*. To knock down the expression of *OsATXR6* (*OsATXR6* RNAi), a 446 bp cDNA fragment, mainly from the 3’UTR region of *OsATXR6* was PCR amplified and cloned into the LH-FAD1390RNAi vector (Zhang et al., 2020).

### *Pst* DC3000 infection and bacteria number counting

*Pst* DC3000 infection was performed as previously described (Yamaguchi et al., 2010). Briefly, *Pst* DC3000 was cultivated with KB medium containing rifampicin at 30°C. After centrifugation, the bacteria were resuspended in sterilized double distilled water (ddH2O) to an OD_600_ of 0.005. The bacterial suspension was infiltrated into *Arabidopsis* leaves using a syringe. Leaf discs with a 6 mm diameter were collected three hours post-inoculation (0 DAI) or three days post-inoculation (3 DAI). The discs were washed with sterilized ddH₂O and ground in 100 μl of sterilized ddH2O. After adding 900 μl of sterilized ddH2O, the samples were thoroughly mixed and serially diluted 1:10. Then, 50 µl of each dilution was plated on TSA agar medium containing rifampicin, and colony numbers were counted after incubating at 30°C for 2 days.

### H_2_O_2_ and NaCl treatment

After-ripened seeds (harvested and stored at room temperature for at least three months) were sown on 1/2 MS medium, with or without H_2_O_2_ or NaCl. After stratification at 4°C for three days, seeds were germinated under long-day conditions at 22°C. Plants with true leaves were counted at 10, 14, or 15 days after germination. Two or three biological replicates were performed for each condition. For the H_2_O_2_ treatment in RT-qPCR and ChIP-qPCR experiments, 3-week-old *Arabidopsis* plants were sprayed with 20mM H_2_O_2_ or with H_2_O (as a mock control). There leaves were collected 5 hours after treatment.

### Measurement of peroxidase activity

Peroxidase activity was measured using the peroxidase activity detection kit (Solarbio, BC0095) following the manufacturer’s instruction. Briefly, 0.1 g of leaves from 3-week-old *Arabidopsis* plants were homogenized on ice using a mortar and pestle in 1 ml extraction buffer. After centrifugation, the supernatant was mixed with buffer 1, 2, and 3, and the absorbance was then measured with a spectrometer at OD_470_.

### Western blotting

Nuclei extracts from leaves of 3-week-old *Arabidopsis* plants or 40-day-old rice were separated by SDS-PAGE and then transferred to a 0.2 μm nitrocellulose membrane (GE Healthcare). Proteins were detected with anti-H3K27me1 (Millipore, 07-448) or anti-H3 (Abcam, ab1791) antibodies. The intensity of the protein bands were quantified using ImageJ and normalized to the H3 loading control.

### DNA content analysis

For DNA content analysis, leaves from 3-week-old *Arabidopsis* plants and 40-day-old rice plants were chopped in Galbraith buffer (20 mM MOPS, 45 mM MgCl2, 30 mM sodium citrate, and 0.1% Triton X-100) for *Arabidopsis* or lysis buffer (25 mM Tris-HCl pH 7.6, 0.44 M sucrose, 10 mM MgCl2, 0.1% Triton X-100, 2 mM spermine, and 10 mM β-Mercaptoethanol) for rice. Then samples were filtered through a 30 μm filter, and nuclei were stained with DAPI. Flow cytometry profiles were obtained using a BD FACSAria II cell sorter.

### Pollen viability examination

Mature rice pollen grains were stained with a 1% I_2_-KI solution and observed under a light microscope to assess viability.

### Immunofluorescence

Immunostaining of isolated leaf nuclei was performed as previously described (Wang et al., 2023a). H3K27me1 signals were detected with anti-H3K27me1 antibody (Millipore, 07-448). Images were captured with a Zeiss confocal laser scanning microscope.

### RNA-seq

For RNA-seq analysis, total RNA was extracted from leaves of 3-week-old *Arabidopsis* plants or 40-day-old rice plants using the Minibest plant RNA extraction kit (Takara, 9769) and three independent biological replicates were performed. Sequencing libraries were prepared with the NEBNext Ultra RNA library prep kit for Illumina (NEB, 7530L) according to the manufacturer’s instruction. Prepared libraries were sequenced on a NovaSeq 6000 platform and paired-end 150bp reads were generated. Adapter trimming was performed and low quality reads were filtered with fastp version 0.20.1 (Chen et al., 2018). Reads were aligned to the *Arabidopsis* genome (TAIR10) or the rice genome (RGAP, version 7.0) using Hisat2 version 2.1.10 (Kim et al., 2019). Reads per gene were counted by HTseq version 0.11.2 (Anders et al., 2015). Transcripts per million (TPM) values were generated using R. Differential gene expression analysis was performed using DESeq2 version 1.26.0 (Love et al., 2014). Genes were considered as differentially expressed if they exhibited a fold change greater than two and a *P* adjust value < 0.05. Gene ontology analysis was conducted with DAVID (https://david.ncifcrf.gov/) (Huang et al., 2009) for *Arabidopsis* and PlantRegMap (https://plantregmap.gao-lab.org/) (Tian et al., 2019) for rice.

### ChIP-seq

ChIP-seq was performed using leaves from 3-week-old *Arabidopsis* plants, which were fixed with 1% formaldehyde. After nuclei extraction, mononucleosomes were generated by digesting with micrococcal nuclease (NEB, M0247S). Immunoprecipitation was conducted with anti-H3K27me3 antibody (Millipore, 07-449) as previously described (Pan et al., 2023). Two independent biological replicates were performed. Immunoprecipitated DNA were subjected to library preparation with the VAHTS universal DNA library prep kit for Illumina (Vazyme, ND607-02) according to the manufacturer’s instruction. Prepared libraries were sequenced on a NovaSeq 6000 platform and paired-end 150bp reads were generated. Adapter trimming was performed and low quality reads were filtered with fastp version 0.20.1 (Chen *et al*., 2018). Reads were mapped to the *Arabidopsis* (TAIR10) genome with Botiew2 version 2.4.2 (Langmead and Salzberg, 2012). Duplicate reads were filtered using Picard version 2.24.0 MarkDuplicates (https://github.com/broadinstitute/picard). H3K27me3 peaks in WT Col were called using MACS2 version 2.1.2 with default parameters (Zhang et al., 2008). Only peaks identified in both biological replicates were retained. For data visualization, data from two biological replicates were merged, and bigwig coverage files were generated using deepTools utility bamCoverage with a bin size of 10bp (Ramirez et al., 2014). Average ChIP-seq profiles were generated using deepTools utility plotProfile.

### RT-qPCR

Total RNA was extracted from leaves of 3-week-old *Arabidopsis* plants or 1-week-old rice seedlings with Minibest plant RNA extraction kit (Takara, 9769) and three independent biological replicates were performed. Reverse transcription was performed using HiScript III 1^st^ Strand cDNA Synthesis Kit (Vazyme, R312-02). Real-time quantitative PCR was conducted on an Applied Biosystems QuantStudio 6 Flex Real-Time PCR System using ChamQ Universal SYBR qPCR Master Mix (Vazyme, Q711-02). For normalization, *TUB2* was used as the endogenous control in *Arabidopsis*, and *OsUBQ5* was used in rice. Primers used for amplification are listed in Supplemental Table 6.

### ChIP-qPCR

Leaves from 3-week-old *Arabidopsis* plants were fixed with 1% formaldehyde. After nuclei extraction, mononucleosomes were generated by digesting with micrococcal nuclease (NEB, M0247S). Immunoprecipitation was conducted with anti-H3K27me1 (Millipore, 07-448), anti-H3K27ac (Abclonal, a7253), or anti-H3K36me3 (Abacm, ab9050) antibody as previously described (Pan *et al*., 2023). The amount of immunoprecipitated DNA was quantified by real-time PCR. Three independent biological replicates were performed. Primers used for amplification are listed in Supplemental Table 6.

## Data availability

The datasets generated in this study are available in the GEO repository under the accession numbers GSE278500 (private access token: glqfqguybbcdzih), GSE278501 (private access token: ybcdqkwobvsjfyh), and GSE278502 (private access token: cvoxwcqudrcnxgr).

## Author contributions

X.L., J.P., and D.J. designed experiments, X.L., J.P., H. L., and D.J. performed experiments, X.L., J.P., H.Z., and Q.L. analyzed data, D.J. wrote the paper with the help from X.L., J.P., and Q.L.

## Acknowledgment

We thank Dr. Lei Li for the valuable advice on the bacterial infection experiment. This work was supported by the National Key R&D Program of China (2023YFD1200704) and the intramural research support from Temasek Life Sciences Laboratory.

## References

Aceituno, F.F., Moseyko, N., Rhee, S.Y., and Gutiérrez, R.A. (2008). The rules of gene expression in plants: Organ identity and gene body methylation are key factors for regulation of gene expression in Arabidopsis thaliana. BMC genomics 9, 438. 10.1186/1471-2164-9-438.

Anders, S., Pyl, P.T., and Huber, W. (2015). HTSeq-a Python framework to work with high-throughput sequencing data. Bioinformatics 31, 166–169. 10.1093/bioinformatics/btu638.

Antunez-Sanchez, J., Naish, M., Ramirez-Prado, J.S., Ohno, S., Huang, Y., Dawson, A., Opassathian, K., Manza-Mianza, D., Ariel, F., Raynaud, C., et al. (2020). A new role for histone demethylases in the maintenance of plant genome integrity. eLife 9. 10.7554/eLife.58533.

Borg, M., Jiang, D., and Berger, F. (2021). Histone variants take center stage in shaping the epigenome. Current opinion in plant biology 61, 101991. 10.1016/j.pbi.2020.101991.

Bourguet, P., Picard, C.L., Yelagandula, R., Pélissier, T., Lorković, Z.J., Feng, S., Pouch-Pélissier, M.-N., Schmücker, A., Jacobsen, S.E., Berger, F., and Mathieu, O. (2021). The histone variant H2A.W and linker histone H1 co-regulate heterochromatin accessibility and DNA methylation. Nature communications 12, 2683. 10.1038/s41467-021-22993-5.

Bowles, A.M.C., Bechtold, U., and Paps, J. (2020). The Origin of Land Plants Is Rooted in Two Bursts of Genomic Novelty. Current biology : CB 30, 530–536.e532. 10.1016/j.cub.2019.11.090.

Buschmann, H., and Holzinger, A. (2020). Understanding the algae to land plant transition. Journal of experimental botany 71, 3241–3246. 10.1093/jxb/eraa196.

Chen, C., Li, C., Wang, Y., Renaud, J., Tian, G., Kambhampati, S., Saatian, B., Nguyen, V., Hannoufa, A., Marsolais, F., et al. (2017). Cytosolic acetyl-CoA promotes histone acetylation predominantly at H3K27 in Arabidopsis. Nature plants 3, 814–824. 10.1038/s41477-017-0023-7.

Chen, S.F., Zhou, Y.Q., Chen, Y.R., and Gu, J. (2018). fastp: an ultra-fast all-in-one FASTQ preprocessor. Bioinformatics 34, 884–890. 10.1093/bioinformatics/bty560.

Clark, J.W. (2023). Genome evolution in plants and the origins of innovation. New Phytologist 240, 2204–2209. 10.1111/nph.19242.

Del Olmo, I., Lopez, J.A., Vazquez, J., Raynaud, C., Pineiro, M., and Jarillo, J.A. (2016). Arabidopsis DNA polymerase recruits components of Polycomb repressor complex to mediate epigenetic gene silencing. Nucleic acids research 44, 5597–5614 10.1093/nar/gkw156.

Ding, Y., Wang, X., Su, L., Zhai, J., Cao, S., Zhang, D., Liu, C., Bi, Y., Qian, Q., Cheng, Z., et al. (2007). SDG714, a histone H3K9 methyltransferase, is involved in Tos17 DNA methylation and transposition in rice. The Plant cell 19, 9–22. 10.1105/tpc.106.048124.

Dong, J., LeBlanc, C., Poulet, A., Mermaz, B., Villarino, G., Webb, K.M., Joly, V., Mendez, J., Voigt, P., and Jacob, Y. (2021). H3.1K27me1 maintains transcriptional silencing and genome stability by preventing GCN5-mediated histone acetylation. The Plant cell 33, 961–979 10.1093/plcell/koaa027.

Du, J., Johnson, L.M., Jacobsen, S.E., and Patel, D.J. (2015). DNA methylation pathways and their crosstalk with histone methylation. Nature reviews. Molecular cell biology 16, 519–532 10.1038/nrm4043.

Ebbs, M.L., Bartee, L., and Bender, J. (2005). H3 lysine 9 methylation is maintained on a transcribed inverted repeat by combined action of SUVH6 and SUVH4 methyltransferases. Molecular and cellular biology 25, 10507–10515. 10.1128/MCB.25.23.10507-10515.2005.

Ebbs, M.L., and Bender, J. (2006). Locus-specific control of DNA methylation by the Arabidopsis SUVH5 histone methyltransferase. The Plant cell 18, 1166–1176. 10.1105/tpc.106.041400.

Ebert, A., Schotta, G., Lein, S., Kubicek, S., Krauss, V., Jenuwein, T., and Reuter, G. (2004). Su(var) genes regulate the balance between euchromatin and heterochromatin in Drosophila. Genes & development 18, 2973–2983. 10.1101/gad.323004.

Eljebbawi, A., Savelli, B., Libourel, C., Estevez, J.M., and Dunand, C. (2022). Class III Peroxidases in Response to Multiple Abiotic Stresses in Arabidopsis thaliana Pyrenean Populations. Int J Mol Sci 23. 10.3390/ijms23073960.

Huang, D.W., Sherman, B.T., and Lempicki, R.A. (2009). Systematic and integrative analysis of large gene lists using DAVID bioinformatics resources. Nature protocols 4, 44–57. 10.1038/nprot.2008.211.

Hyun, Y., Yun, H., Park, K., Ohr, H., Lee, O., Kim, D.H., Sung, S., and Choi, Y. (2013). The catalytic subunit of Arabidopsis DNA polymerase alpha ensures stable maintenance of histone modification. Development 140, 156–166. 10.1242/dev.084624.

Jackson, J.P., Lindroth, A.M., Cao, X., and Jacobsen, S.E. (2002). Control of CpNpG DNA methylation by the KRYPTONITE histone H3 methyltransferase. Nature 416, 556–560. 10.1038/nature731.

Jacob, Y., Bergamin, E., Donoghue, M.T., Mongeon, V., LeBlanc, C., Voigt, P., Underwood, C.J., Brunzelle, J.S., Michaels, S.D., Reinberg, D., et al. (2014). Selective methylation of histone H3 variant H3.1 regulates heterochromatin replication. Science 343, 1249–1253. 10.1126/science.1248357.

Jacob, Y., Feng, S., LeBlanc, C.A., Bernatavichute, Y.V., Stroud, H., Cokus, S., Johnson, L.M., Pellegrini, M., Jacobsen, S.E., and Michaels, S.D. (2009). ATXR5 and ATXR6 are H3K27 monomethyltransferases required for chromatin structure and gene silencing. Nature structural & molecular biology 16, 763–768. 10.1038/nsmb.1611.

Jacob, Y., and Michaels, S.D. (2009). H3K27me1 is E(z) in animals, but not in plants. Epigenetics 4, 366–369.

Jacob, Y., Stroud, H., Leblanc, C., Feng, S., Zhuo, L., Caro, E., Hassel, C., Gutierrez, C., Michaels, S.D., and Jacobsen, S.E. (2010). Regulation of heterochromatic DNA replication by histone H3 lysine 27 methyltransferases. Nature 466, 987–991. 10.1038/nature09290.

Jiang, D., and Berger, F. (2017). DNA replication-coupled histone modification maintains Polycomb gene silencing in plants. Science 357, 1146–1149. 10.1126/science.aan4965.

Karimi, M., Inze, D., and Depicker, A. (2002). GATEWAY vectors for Agrobacterium-mediated plant transformation. Trends in plant science 7, 193–195. 10.1016/s1360-1385(02)02251-3.

Kidwai, M., Ahmad, I.Z., and Chakrabarty, D. (2020). Class III peroxidase: an indispensable enzyme for biotic/abiotic stress tolerance and a potent candidate for crop improvement. Plant cell reports 39, 1381–1393. 10.1007/s00299-020-02588-y.

Kim, D., Paggi, J.M., Park, C., Bennett, C., and Salzberg, S.L. (2019). Graph-based genome alignment and genotyping with HISAT2 and HISAT-genotype. Nat Biotechnol 37, 907-+. 10.1038/s41587-019-0201-4.

Kim, S.Y., He, Y.H., Jacob, Y., Noh, Y.S., Michaels, S., and Amasino, R. (2005). Establishment of the vernalization-responsive, winter-annual habit in Arabidopsis requires a putative histone H3 methyl transferase. The Plant cell 17, 3301–3310. 10.1105/tpc.105.034645.

Kornet, N., and Scheres, B. (2009). Members of the GCN5 histone acetyltransferase complex regulate PLETHORA-mediated root stem cell niche maintenance and transit amplifying cell proliferation in Arabidopsis. The Plant cell 21, 1070–1079. 10.1105/tpc.108.065300.

Kumar, P., Kumar, P., Verma, V., Irfan, M., Sharma, R., and Bhargava, B. (2023). How plants conquered land: evolution of terrestrial adaptation. Journal of Evolutionary Biology 35, 5–14. 10.1111/jeb.14062.

Langmead, B., and Salzberg, S.L. (2012). Fast gapped-read alignment with Bowtie 2. Nature methods 9, 357–U354. 10.1038/Nmeth.1923.

Love, M.I., Huber, W., and Anders, S. (2014). Moderated estimation of fold change and dispersion for RNA-seq data with DESeq2. Genome biology 15. ARTN 550 10.1186/s13059-014-0550-8.

Lu, F., Cui, X., Zhang, S., Jenuwein, T., and Cao, X. (2011). Arabidopsis REF6 is a histone H3 lysine 27 demethylase. Nature genetics 43, 715–719. 10.1038/ng.854.

Ma, X., Zhang, Q., Zhu, Q., Liu, W., Chen, Y., Qiu, R., Wang, B., Yang, Z., Li, H., Lin, Y., et al. (2015). A Robust CRISPR/Cas9 System for Convenient, High-Efficiency Multiplex Genome Editing in Monocot and Dicot Plants. Molecular plant 8, 1274–1284. 10.1016/j.molp.2015.04.007.

Mathieu, O., Probst, A.V., and Paszkowski, J. (2005). Distinct regulation of histone H3 methylation at lysines 27 and 9 by CpG methylation in Arabidopsis. The EMBO journal 24, 2783–2791. 10.1038/sj.emboj.7600743.

Mbadinga Mbadinga, D.L., Li, Q., Ranocha, P., Martinez, Y., and Dunand, C. (2020). Global analysis of non-animal peroxidases provides insights into the evolution of this gene family in the green lineage. Journal of experimental botany 71, 3350–3360. 10.1093/jxb/eraa141.

Montgomery, S.A., Tanizawa, Y., Galik, B., Wang, N., Ito, T., Mochizuki, T., Akimcheva, S., Bowman, J.L., Cognat, V., Marechal-Drouard, L., et al. (2020). Chromatin Organization in Early Land Plants Reveals an Ancestral Association between H3K27me3, Transposons, and Constitutive Heterochromatin. Current biology : CB 30, 573–588 e577. 10.1016/j.cub.2019.12.015.

Naumann, K., Fischer, A., Hofmann, I., Krauss, V., Phalke, S., Irmler, K., Hause, G., Aurich, A.C., Dorn, R., Jenuwein, T., and Reuter, G. (2005). Pivotal role of AtSUVH2 in heterochromatic histone methylation and gene silencing in Arabidopsis. The EMBO journal 24, 1418–1429. 10.1038/sj.emboj.7600604.

Pan, J., Zhang, H., Zhan, Z., Zhao, T., and Jiang, D. (2023). A REF6-dependent H3K27me3-depleted state facilitates gene activation during germination in Arabidopsis. Journal of Genetics and Genomics 50, 178–191. 10.1016/j.jgg.2022.09.001.

Passardi, F., Bakalovic, N., Teixeira, F.K., Margis-Pinheiro, M., Penel, C., and Dunand, C. (2007). Prokaryotic origins of the non-animal peroxidase superfamily and organelle-mediated transmission to eukaryotes. Genomics 89, 567–579. 10.1016/j.ygeno.2007.01.006.

Passardi, F., Longet, D., Penel, C., and Dunand, C. (2004). The class III peroxidase multigenic family in rice and its evolution in land plants. Phytochemistry 65, 1879–1893. 10.1016/j.phytochem.2004.06.023.

Pedreira, J., Herrera, M.T., Zarra, I., and Revilla, G. (2011). The overexpression of AtPrx37, an apoplastic peroxidase, reduces growth in Arabidopsis. Physiol Plant 141, 177–187. 10.1111/j.1399-3054.2010.01427.x.

Potok, M.E., Zhong, Z., Picard, C.L., Liu, Q., Do, T., Jacobsen, C.E., Sakr, O., Naranbaatar, B., Thilakaratne, R., Khnkoyan, Z., et al. (2022). The role of ATXR6 expression in modulating genome stability and transposable element repression in Arabidopsis. Proceedings of the National Academy of Sciences of the United States of America 119. 10.1073/pnas.2115570119.

Raggi, S., Ferrarini, A., Delledonne, M., Dunand, C., Ranocha, P., De Lorenzo, G., Cervone, F., and Ferrari, S. (2015). The Arabidopsis Class III Peroxidase AtPRX71 Negatively Regulates Growth under Physiological Conditions and in Response to Cell Wall Damage. Plant physiology 169, 2513–2525. 10.1104/pp.15.01464.

Ramirez, F., Dundar, F., Diehl, S., Gruning, B.A., and Manke, T. (2014). deepTools: a flexible platform for exploring deep-sequencing data. Nucleic acids research 42, W187–W191 10.1093/nar/gku365.

Raynaud, C., Sozzani, R., Glab, N., Domenichini, S., Perennes, C., Cella, R., Kondorosi, E., and Bergounioux, C. (2006). Two cell-cycle regulated SET-domain proteins interact with proliferating cell nuclear antigen (PCNA) in Arabidopsis. The Plant journal : for cell and molecular biology 47, 395–407. 10.1111/j.1365-313X.2006.02799.x.

Schonrock, N., Bouveret, R., Leroy, O., Borghi, L., Kohler, C., Gruissem, W., and Hennig, L. (2006). Polycomb-group proteins repress the floral activator AGL19 in the FLC-independent vernalization pathway. Genes & development 20, 1667–1678. 10.1101/gad.377206.

Shen, X., Liu, Y., Hsu, Y.J., Fujiwara, Y., Kim, J., Mao, X., Yuan, G.C., and Orkin, S.H. (2008). EZH1 mediates methylation on histone H3 lysine 27 and complements EZH2 in maintaining stem cell identity and executing pluripotency. Molecular cell 32, 491–502. 10.1016/j.molcel.2008.10.016.

Stroud, H., Hale, C.J., Feng, S., Caro, E., Jacob, Y., Michaels, S.D., and Jacobsen, S.E. (2012). DNA methyltransferases are required to induce heterochromatic re-replication in Arabidopsis. PLoS genetics 8, e1002808. 10.1371/journal.pgen.1002808.

Tian, F., Yang, D.-C., Meng, Y.-Q., Jin, J., and Gao, G. (2019). PlantRegMap: charting functional regulatory maps in plants. Nucleic acids research 48, D1104–D1113 10.1093/nar/gkz1020.

Vaillant, I., and Paszkowski, J. (2007). Role of histone and DNA methylation in gene regulation. Current opinion in plant biology 10, 528–533. 10.1016/j.pbi.2007.06.008.

Vongs, A., Kakutani, T., Martienssen, R.A., and Richards, E.J. (1993). Arabidopsis thaliana DNA methylation mutants. Science 260, 1926–1928. 10.1126/science.8316832.

Wang, L., Xue, M., Zhang, H., Ma, L., and Jiang, D. (2023a). TONSOKU is required for the maintenance of repressive chromatin modifications in Arabidopsis. Cell reports 42, 112738. 10.1016/j.celrep.2023.112738.

Wang, L., Xue, M., Zhang, H., Ma, L., and Jiang, D. (2023b). TONSOKU is required for the maintenance of repressive chromatin modifications in Arabidopsis. Cell reports 42, 112738. 10.1016/j.celrep.2023.112738.

Wang, S., Durrant, W.E., Song, J., Spivey, N.W., and Dong, X. (2010). Arabidopsis BRCA2 and RAD51 proteins are specifically involved in defense gene transcription during plant immune responses. Proceedings of the National Academy of Sciences of the United States of America 107, 22716–22721. 10.1073/pnas.1005978107.

Wiles, E.T., and Selker, E.U. (2017). H3K27 methylation: a promiscuous repressive chromatin mark. Curr Opin Genet Dev 43, 31–37. 10.1016/j.gde.2016.11.001.

Wodniok, S., Brinkmann, H., Glöckner, G., Heidel, A.J., Philippe, H., Melkonian, M., and Becker, B. (2011). Origin of land plants: Do conjugating green algae hold the key? BMC Evolutionary Biology 11, 104. 10.1186/1471-2148-11-104.

Xu, L., Zhao, Z., Dong, A., Soubigou-Taconnat, L., Renou, J.P., Steinmetz, A., and Shen, W.H. (2008). Di- and tri- but not monomethylation on histone H3 lysine 36 marks active transcription of genes involved in flowering time regulation and other processes in Arabidopsis thaliana. Molecular and cellular biology 28, 1348–1360. 10.1128/MCB.01607-07.

Yamaguchi, Y., Huffaker, A., Bryan, A.C., Tax, F.E., and Ryan, C.A. (2010). PEPR2 is a second receptor for the Pep1 and Pep2 peptides and contributes to defense responses in Arabidopsis. The Plant cell 22, 508–522. 10.1105/tpc.109.068874.

Yan, S., Wang, W., Marqués, J., Mohan, R., Saleh, A., Durrant, Wendy E., Song, J., and Dong, X. (2013). Salicylic Acid Activates DNA Damage Responses to Potentiate Plant Immunity. Molecular cell 52, 602–610. 10.1016/j.molcel.2013.09.019.

Yu, C.W., Tai, R., Wang, S.C., Yang, P., Luo, M., Yang, S., Cheng, K., Wang, W.C., Cheng, Y.S., and Wu, K. (2017). HISTONE DEACETYLASE6 Acts in Concert with Histone Methyltransferases SUVH4, SUVH5, and SUVH6 to Regulate Transposon Silencing. The Plant cell 29, 1970–1983. 10.1105/tpc.16.00570.

Zhang, B.Q., Liu, X.S., Feng, S.J., Zhao, Y.N., Wang, L.L., Rono, J.K., Li, H., and Yang, Z.M. (2020). Developing a cadmium resistant rice genotype with OsHIPP29 locus for limiting cadmium accumulation in the paddy crop. Chemosphere 247, 125958. 10.1016/j.chemosphere.2020.125958.

Zhang, X., Clarenz, O., Cokus, S., Bernatavichute, Y.V., Pellegrini, M., Goodrich, J., and Jacobsen, S.E. (2007). Whole-genome analysis of histone H3 lysine 27 trimethylation in Arabidopsis. PLoS biology 5, e129. 10.1371/journal.pbio.0050129.

Zhang, Y., Liu, T., Meyer, C.A., Eeckhoute, J., Johnson, D.S., Bernstein, B.E., Nusbaum, C., Myers, R.M., Brown, M., Li, W., and Liu, X.S. (2008). Model-based analysis of ChIP-Seq (MACS). Genome biology 9, R137. 10.1186/gb-2008-9-9-r137.

Zhao, T., Zhan, Z.P., and Jiang, D.H. (2019). Histone modifications and their regulatory roles in plant development and environmental memory. Journal of Genetics and Genomics 46, 467–476. 10.1016/j.jgg.2019.09.005.

Zhao, X., Wang, J., Jin, D., Cheng, J., Chen, H., Li, Z., Wang, Y., Lou, H., Zhu, J.-K., Du, X., and Gong, Z. (2023). AtMCM10 promotes DNA replication-coupled nucleosome assembly in Arabidopsis. J Integr Plant Biol 65, 203–222. 10.1111/jipb.13438.

Zheng, S., Hu, H., Ren, H., Yang, Z., Qiu, Q., Qi, W., Liu, X., Chen, X., Cui, X., Li, S., et al. (2019). The Arabidopsis H3K27me3 demethylase JUMONJI 13 is a temperature and photoperiod dependent flowering repressor. Nature communications 10, 1303. 10.1038/s41467-019-09310-x.

Zhou, Y., Tergemina, E., Cui, H., Forderer, A., Hartwig, B., Velikkakam James, G., Schneeberger, K., and Turck, F. (2017). Ctf4-related protein recruits LHP1-PRC2 to maintain H3K27me3 levels in dividing cells in Arabidopsis thaliana. Proceedings of the National Academy of Sciences of the United States of America 114, 4833–4838. 10.1073/pnas.1620955114.

