## Supplemental Figures and Tables for "Land plant-specific H3K27 methyltransferases ATXR5 and ATXR6 control plant development and stress responses"

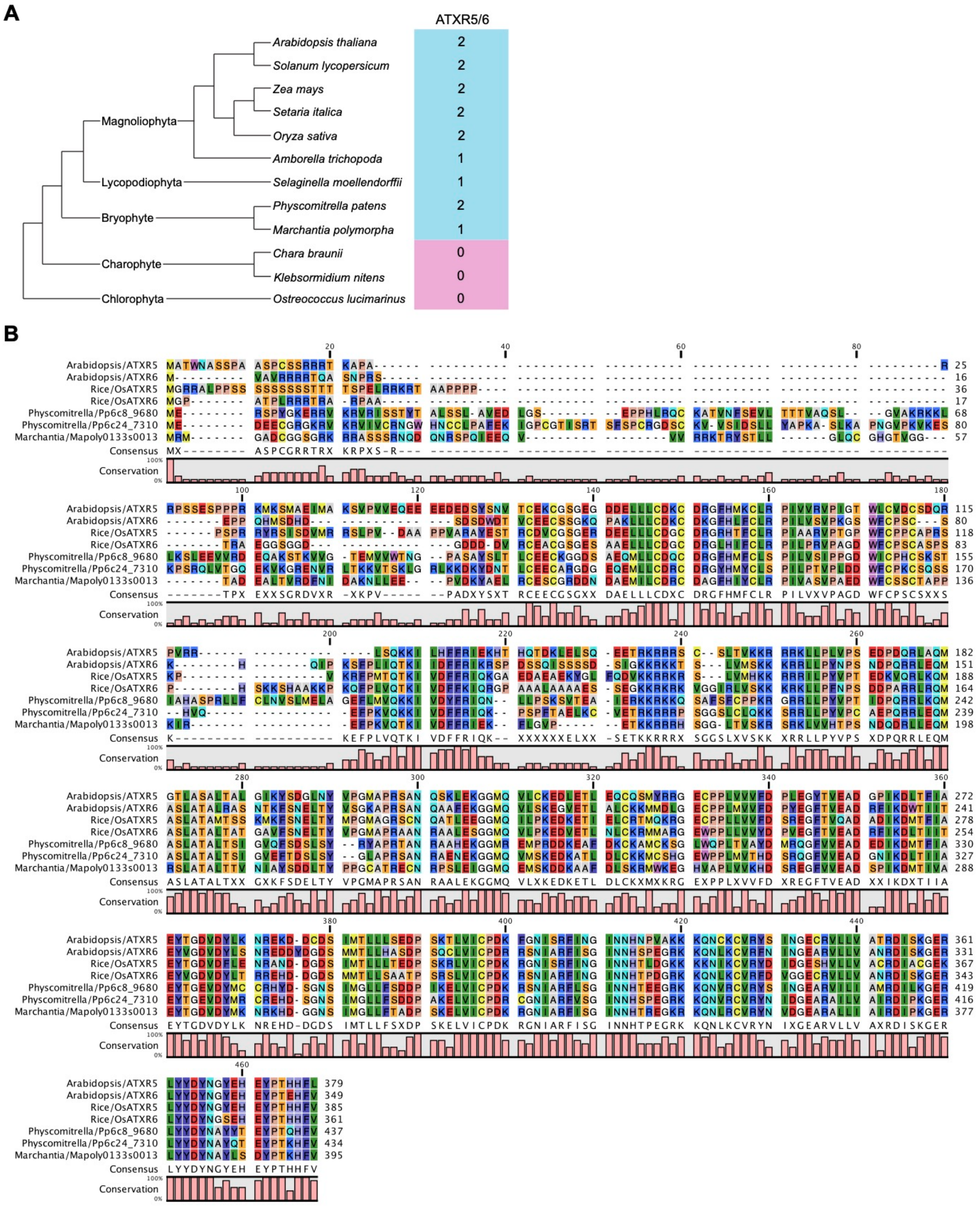


**Supplemental Figure 1. Evolutionary analysis of ATXR5/6 in the plant lineage**

**A.** Number of ATXR5/6 orthologs in representative species across different groups. The blue shading indicates the presence of ATXR5/6, while the pink shading indicates the absence of proteins homologous to ATXR5/6.

**B.** Sequence alignment of protein homologues to ATXR5/6 in *Arabidopsis thaliana*, *Oryza sativa*, *Physcomitrella patens*, *Marchantia polymorpha*.

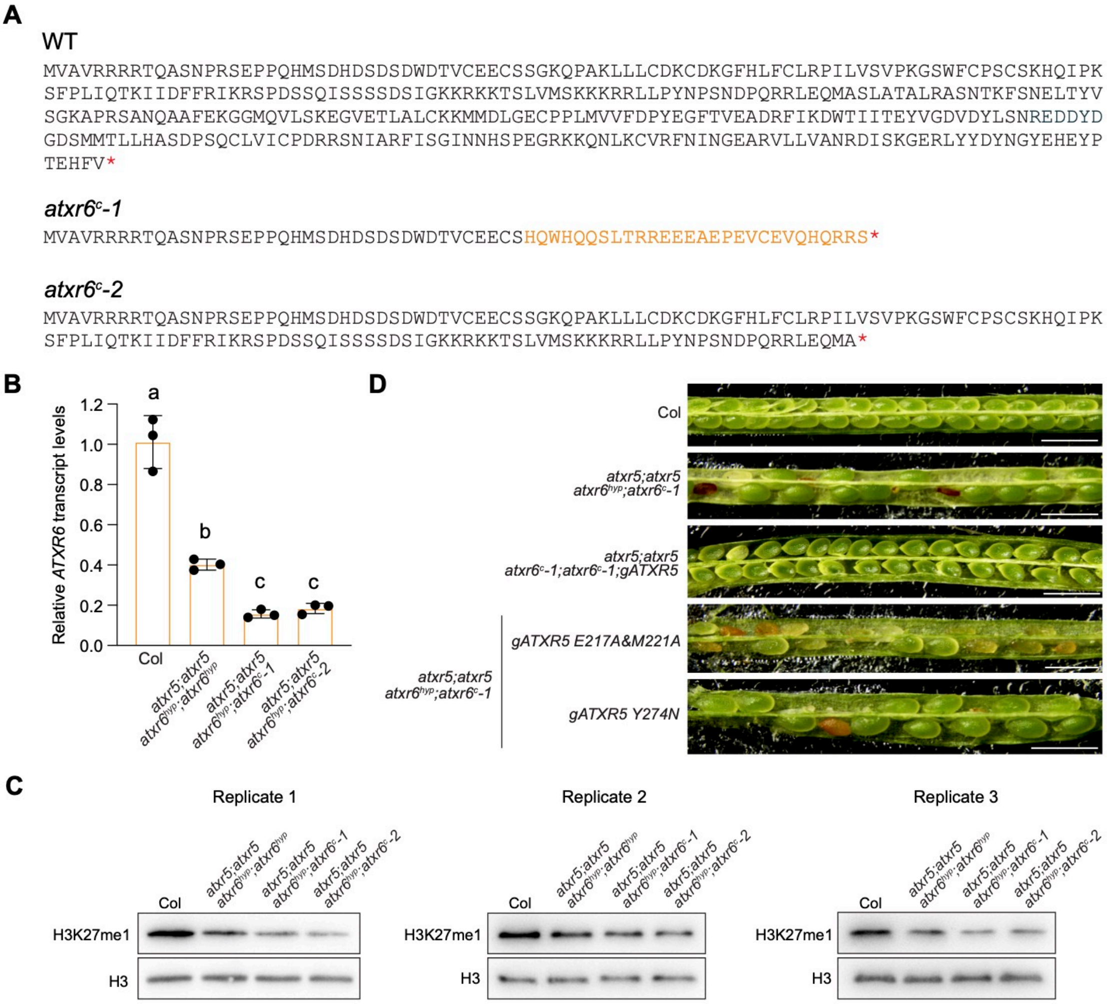


**Supplemental Figure 2. Analyses of molecular and phenotypic changes following the loss of ATXR5 and ATXR6**

**A.** ATXR6 protein sequences in WT Col, *axtr6*$^{c}$*-1*, and *atxr6*$^{c}$*-2*. Mutations in *axtr6*$^{c}$*-1* and *atxr6*$^{c}$*-2* result in frameshift mutations and early termination.

**B.** Relative *ATXR6* transcript levels in the indicated lines determined by RT-qPCR. Values are means ± SD of three biological replicates. The significance of differences was tested using one-way ANOVA with Tukey's test ($P$ < 0.05), with different letters indicating statistically significant differences.

**C.** H3K27me1 levels in in the indicated lines determined by western blotting. H3 was employed as a loading control. Three biological replicates are presented.

**D.** Seed developmental phenotypes of the complementation lines. Scale bar, 1mm.

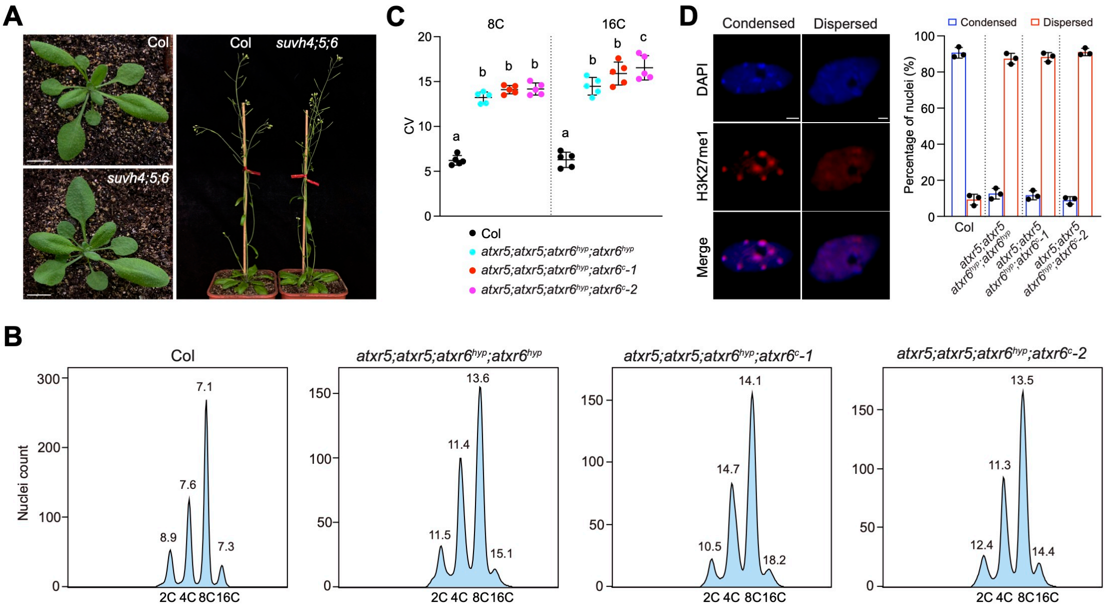


**Supplemental Figure 3. Comparison of heterochromatin defects in weak and strong *atxr5;atxr6* mutants**

**A.** Plant growth phenotypes of Col and the *suvh4;5;6* mutant. Scale bars, 1cm.

**B.** Flow cytometry profiles of leaf nuclei for the indicated lines. The numbers below the peaks indicate the ploidy levels of the nuclei, while the numbers above indicate the coefficient of variation (CV).

**C.** CV values of 8C and 16C nuclei obtained from flow cytometry analysis in the indicated lines. Values are means ± SD of five biological replicates. The significance of differences was tested using one-way ANOVA with Tukey's test ($P < 0.05$), with different letters indicating statistically significant differences.

**D.** Condensed and decondensed chromocenters observed in leaf nuclei stained with DAPI and immunostained with H3K27me1. The bar chart represents the percentages of nuclei showing condensed and dispersed chromocenters in the indicated lines. Values are means ± SD of three biological replicates. At least 100 nuclei were analysed in each replicate. Scale bars, 2 um.

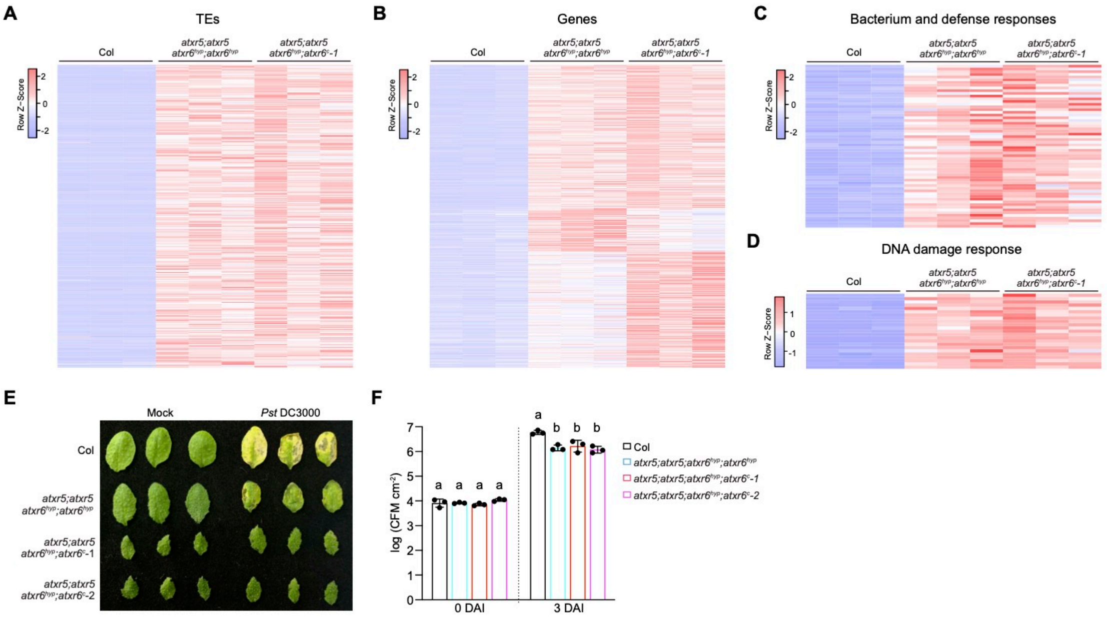


**Supplemental Figure 4. Gene expression and bacterial infection analysis in *atxr5;atxr6* mutants**

**A and B.** Heatmaps showing transcript level significantly increased TEs (A) and genes (B) in $atxr5;atxr6^{hyp}$ and $atxr5;atxr5;atxr6^{hyp};atxr6^{c}$*-1* determined by RNA-seq. Results from three biological replicates are shown.

**C and D.** Heatmaps showing transcript levels of bacterium response (C) and DNA damage response (D) genes determined by RNA-seq. Results from three biological replicates are shown.

**E.** Leaf phenotypes of the indicated lines with or without *Pst* DC3000 infection.

**F.** Bacterial growth assay of *Pst* DC3000 inoculated on the indicated lines. Growth was measured at 0 and 3 days after inoculation (DAI). Values are means ± SD of three biological replicates. The significance of differences was tested using one-way ANOVA with Tukey's test ($P < 0.05$), with different letters indicating statistically significant differences.

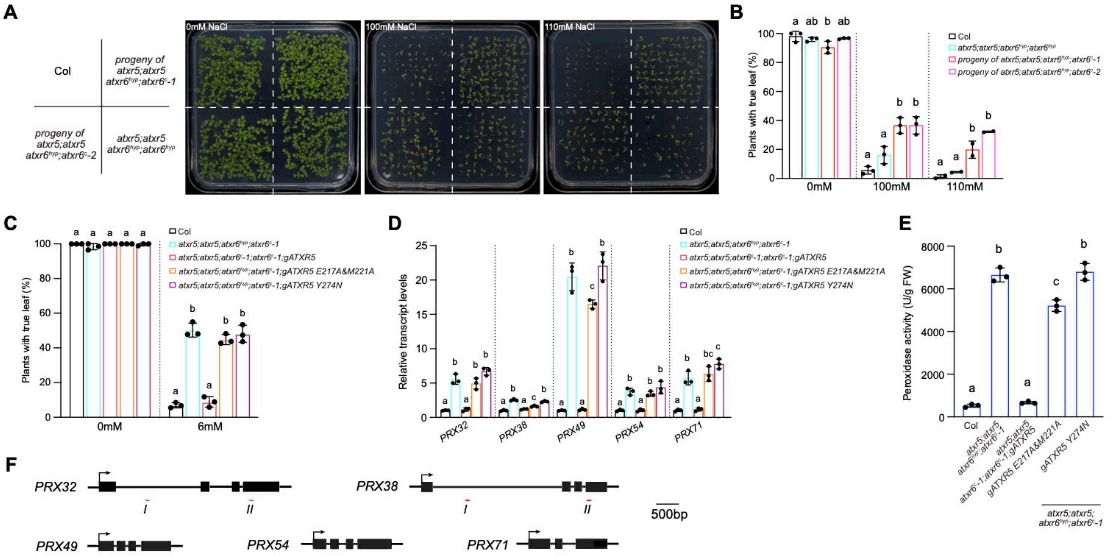


**Supplemental Figure 5. Oxidative and salt stress responses in *atxr5;atxr6* mutants**

**A.** Seedling growth phenotypes on 1/2MS plate supplemented without NaCl or with 100mM or 110mM NaCl. Pictures were taken 10 days after germination.

**B.** True leaf formation rates of seedlings germinated on 1/2MS plate supplemented without NaCl or with 100mM or 110mM NaCl. Rates were measured 14 days after germination. Values are means ± SD of two or three biological replicates. At least 79 seeds were sown for each replicate. The significance of differences was tested using one-way ANOVA with Tukey's test ($P$ < 0.05), with different letters indicating statistically significant differences.

**C.** True leaf formation rates of the complementation lines germinated on 1/2MS plate supplemented without $H_2O_2$ or with 6mM $H_2O_2$. Rates were measured 10 days after germination. Values are means ± SD of three biological replicates. At least 69 seeds were sown for each replicate. The significance of differences was tested using one-way ANOVA with Tukey's test ($P$ < 0.05), with different letters indicating statistically significant differences.

**D.** Transcript levels of peroxidase-coding genes in the complementation lines determined by RT-qPCR. Values are means ± SD of three biological replicates. The significance of differences was tested using one-way ANOVA with Tukey's test ($P$ < 0.05), with different letters indicating statistically significant differences.

**E.** Peroxidase activity in the complementation lines. Values are means ± SD of three biological replicates. The significance of differences was tested using one-way ANOVA with Tukey's test ($P$ < 0.05), with different letters indicating statistically significant differences.

**F.** Schematic structures of *PRX32*, *PRX38*, *PRX49*, *PRX54*, and *PRX71*. Arrows indicate transcription start sites, and red lines indicate regions examined by ChIP-qPCR.

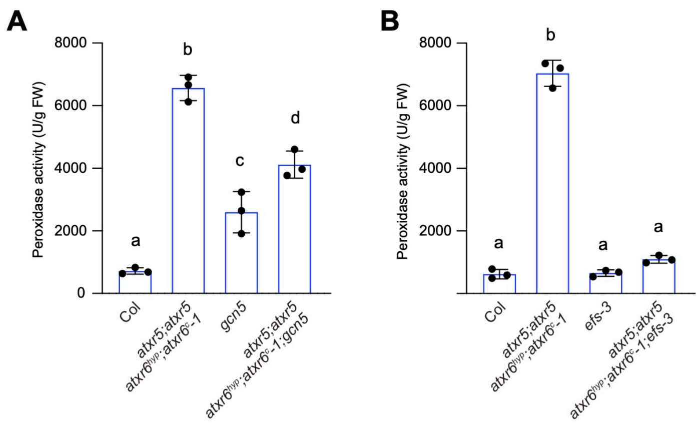


**Supplemental Figure 6. Peroxidase activity following loss of GCN5 (A) or EFS (B) in *atxr5;atxr5;atxr6*$^{hyp}$*;atxr6*$^{c}$*-1***

Values are means ± SD of three biological replicates. The significance of differences was tested using one-way ANOVA with Tukey’s test ($P < 0.05$), with different letters indicating statistically significant differences.

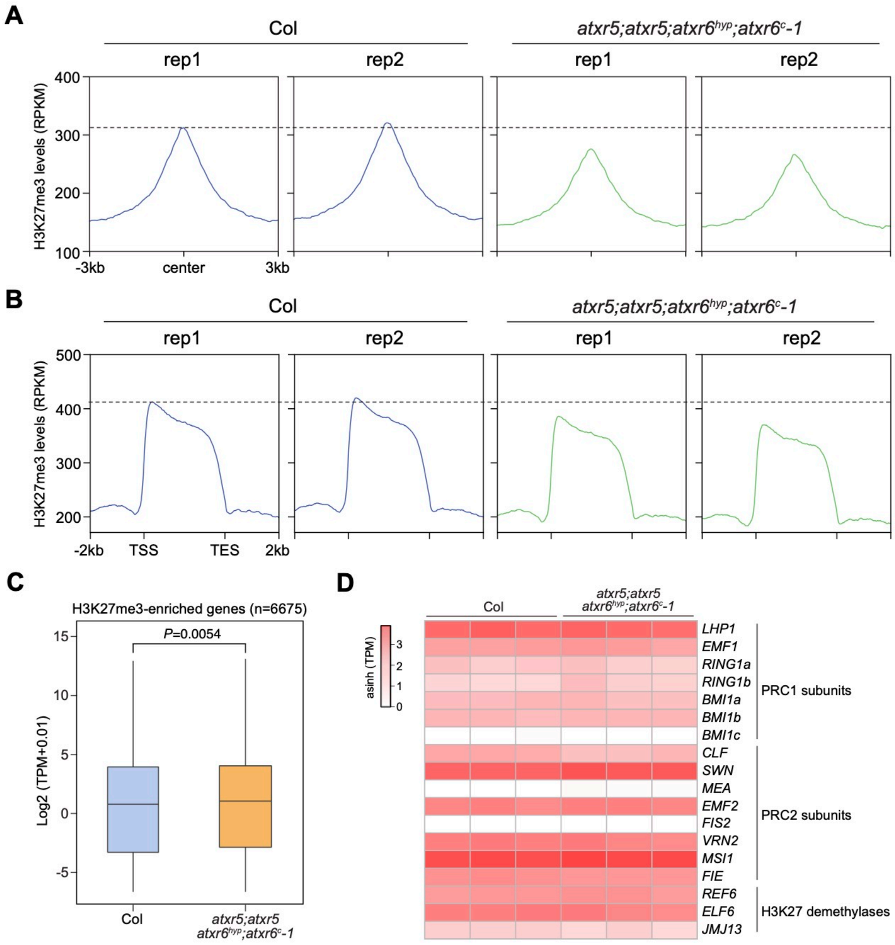


**Supplemental Figure 7. H3K27me3 analysis in Col and *atxr5;atxr5;atxr6$^{hyp}$;atxr6$^{c}$-1***

**A and B.** Metaplots of H3K27me3 ChIP-seq signals in Col and *atxr5;atxr5;atxr6$^{hyp}$;atxr6$^{c}$-1* over H3K27me3-enriched peaks (A) and genes (B) in WT Col. Results from two biological replicates are shown.

**C.** Overall gene expression profiles of H3K27me3-enriched genes in Col and *atxr5;atxr5;atxr6$^{hyp}$;atxr6$^{c}$-1*. The expression values represent the average of three biological replicates. *P* value is based on the Manne-Whitney U test.

**D.** Heatmap showing transcript levels of PcG genes and H3K27 demethylase-coding genes determined by RNA-seq. Results from three biological replicates are shown. PRC1/2 refers to Polycomb group complex 1/2.

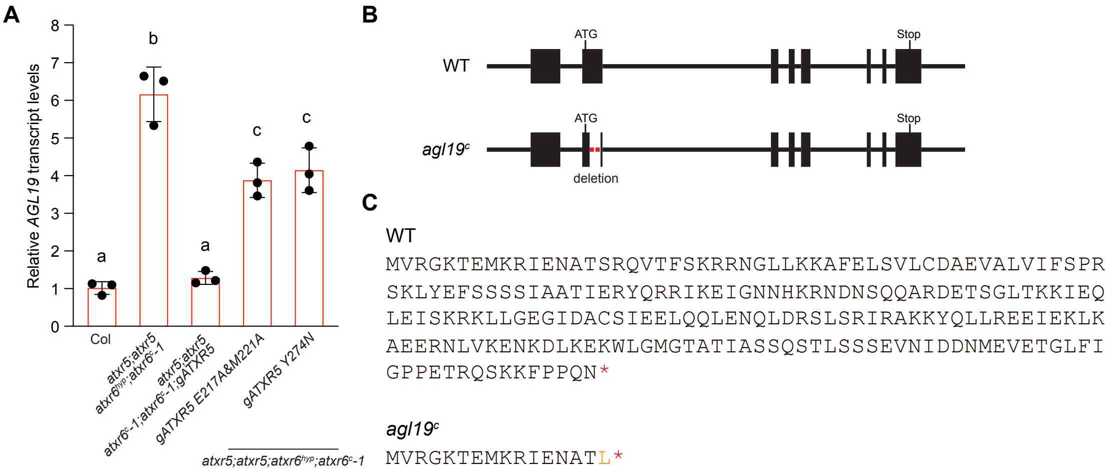


**Supplemental Figure 8. Analysis of *AGL19* expression and CRISPR/Cas9 editing of *AGL19***

**A.** Relative transcript levels of *AGL19* in the complementation lines determined by RT-qPCR. Values are means ± SD of three biological replicates. The significance of differences was tested using one-way ANOVA with Tukey's test ($P < 0.05$), with different letters indicating statistically significant differences.

**B.** Schematic view of the full-length *AGL19* genomic structure. Filled boxes indicate exons, and the red dashed line represents the deleted region in *agl19*$^{c}$.

**C.** AGL19 protein sequences in WT Col and *agl19*$^{c}$. Mutations in *agl19*$^{c}$ results in frameshift mutations and early termination.

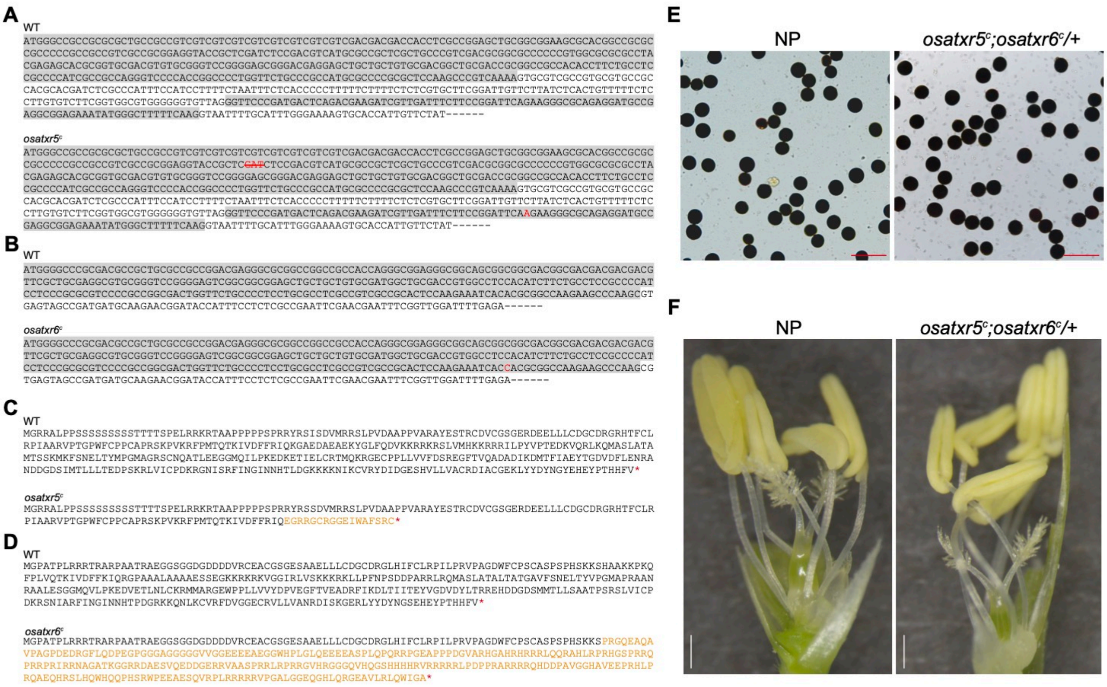


**Supplemental Figure 9. Analyses of molecular and phenotypic changes following loss of OsATXR5 and OsATXR6**

**A and B.** DNA sequences of *OsATXR5* (A) and *OsATXR6* (B) in WT NP, *osatxr5*$^{c}$, and *osatxr6*$^{c}$. Red fonts represent deleted or inserted nucleotides in *osatxr5*$^{c}$ and *osatxr6*$^{c}$.

**C and D.** Protein sequences of OsATXR5 (a) and OsATXR6 (b) in WT NP, *osatxr5*$^{c}$, and *osatxr6*$^{c}$. Mutations in *osatxr5*$^{c}$ and *osatxr6*$^{c}$ result in frameshift mutation and early termination.

**E.** I2-KI staining of pollen from NP and *osatxr5*$^{c}$;*osatxr6*$^{c}$/+. Scale bars, 100um.

**F.** Flower phenotypes of NP and *osatxr5*$^{c}$;*osatxr6*$^{c}$/+. Both lemma and palea have been removed to show the inner floral organs. Scale bars, 0.5mm.

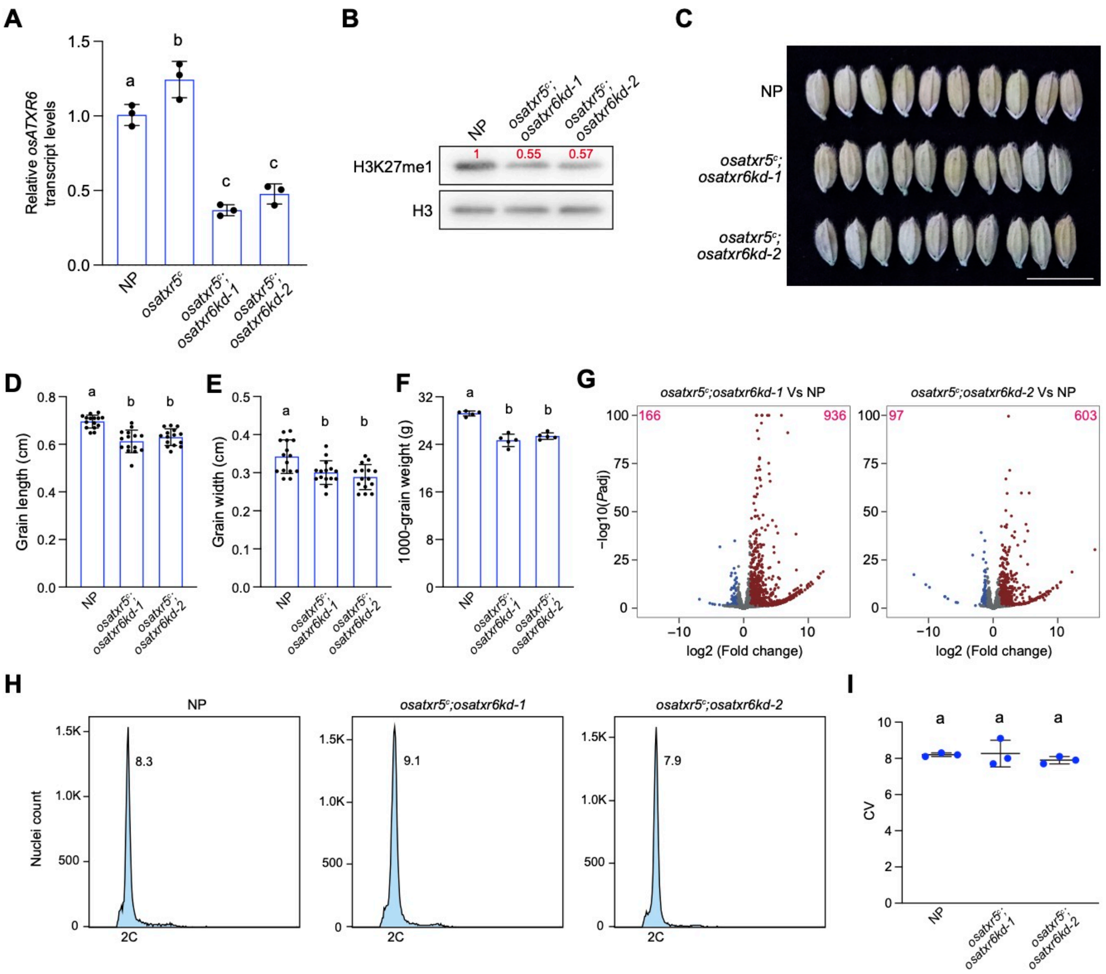


**Supplemental Figure 10. Gene expression and DNA content analysis in *osatxr5c;osatxr6kd* lines**

**A.** Relative Os*ATXR6* transcript levels in the indicated lines determined by RT-qPCR. Values are means ± SD of three biological replicates. The significance of differences was tested using one-way ANOVA with Tukey's test ($P < 0.05$), with different letters indicating statistically significant differences.

**B.** H3K27me1 levels in NP, *osatxr5c;osatxr6kd-1*, and *osatxr5c;osatxr6kd-2* determined by western blotting. H3 was employed as a loading control. Values are fold changes over NP.

**C.** Grain phenotypes of NP, *osatxr5c;osatxr6kd-1*, and *osatxr5c;osatxr6kd-2*. Scale bar, 1cm.

**D-F.** Grain length (D), grain width (E), and 1000-grain weight (F) of NP, *osatxr5c;osatxr6kd-1*, and *osatxr5c;osatxr6kd-2*. 15 grains were measured for grain length and width, and 5 biological replicates were performed for 1000-

grain weight measurements. Values are means ± SD. The significance of differences was tested using one-way ANOVA with Tukey's test ($P$ < 0.05), with different letters indicating statistically significant differences.

**G.** Volcano plots of differentially expressed TEs. The y-axis represents -log10 ($P$ adjust), and the x-axis indicates log2 (fold change). TEs exhibiting at least a two-fold change in expression and an adjusted $P$ value of less than 0.05 are considered misexpressed. The numbers of TEs with increased and decreased transcript levels are indicated in the top right and left corners, respectively.

**H.** Flow cytometry profiles of leaf nuclei from NP, *osatxr5$^{c}$;osatxr6kd-1*, and *osatxr5$^{c}$;osatxr6kd-2*. Only 2C nuclei were detected. The numbers indicate the coefficient of variation (CV).

**I.** CV values of 2C nuclei obtained from flow cytometry analysis in NP, *osatxr5$^{c}$;osatxr6kd-1*, and *osatxr5$^{c}$;osatxr6kd-2*. Values are means ± SD of three biological replicates. The significance of differences was tested using one-way ANOVA with Tukey's test ($P$ < 0.05), with different letters indicating statistically significant differences.

**Supplemental Table 1.** ***Arabidopsis*** **Progenies produced from self-pollination**

| Parent | Progeny | | |
|---|---|---|---|
| *atxr5;atxr5 atxr6*$^{hyp}$*;atxr6*$^{c}$*-1* | *atxr5;atxr5 atxr6*$^{hyp}$*;atxr6*$^{hyp}$ | *atxr5;atxr5 atxr6*$^{hyp}$*;atxr6*$^{c}$*-1* | *atxr5;atxr5 atxr6*$^{c}$*-1;atxr6*$^{c}$*-1* |
| | 81 | 91 | 0 |
| *atxr5;atxr5 atxr6*$^{hyp}$*;atxr6*$^{c}$*-2* | *atxr5;atxr5 atxr6*$^{hyp}$*;atxr6*$^{hyp}$ | *atxr5;atxr5 atxr6*$^{hyp}$*;atxr6*$^{c}$*-2* | *atxr5;atxr5 atxr6*$^{c}$*-2;atxr6*$^{c}$*-2* |
| | 60 | 62 | 0 |

**Supplemental Table 2.** ***Arabidopsis*** **progenies produced from reciprocal crosses**

| Parents (female x male) | Progeny | |
|---|---|---|
| | *atxr5/+;atxr6*$^{hyp}$*/+* | *atxr5/+;atxr6*$^{c}$*/+* |
| *atxr5;atxr5;atxr6*$^{hyp}$*;atxr6*$^{c}$*-1* x WT | 66 | 6 |
| WT x *atxr5;atxr5;atxr6*$^{hyp}$*;atxr6*$^{c}$*-1* | 40 | 42 |
| *atxr5;atxr5;atxr6*$^{hyp}$*;atxr6*$^{c}$*-2* x WT | 86 | 3 |
| WT x *atxr5;atxr5;atxr6*$^{hyp}$*;atxr6*$^{c}$*-2* | 56 | 40 |

**Supplemental Table 3. Rice Progenies produced from self-pollination**

| Parent | Progeny | | |
|---|---|---|---|
| *osatxr5*$^{c}$*; osatxr6*$^{c}$*/+* | *osatxr5*$^{c}$ | *osatxr5*$^{c}$*; osatxr6*$^{c}$*/+* | *osatxr5c; osatxr6*$^{c}$ |
| | 45 | 96 | 0 |

**Supplemental Table 4. Rice progenies produced from reciprocal crosses**

| Parents (female x male) | Progeny | |
|---|---|---|
| | *osatxr5*$^{c}$*/+* | *osatxr5*$^{c}$*/+;osatxr6*$^{c}$*/+* |
| *osatxr5*$^{c}$*;osatxr6*$^{c}$*/+* x WT | 13 | 18 |
| WT x *osatxr5*$^{c}$*;osatxr6*$^{c}$*/+* | 17 | 16 |

**Supplemental Table 5. GuideRNA sequences**

| Mutant | Guide RNA |
|---|---|
| *atxr6$^c$-1* | GCGATGTTACTGCGTCTGTC |
| | GCTAGAGCAAATGGCGTCTC |
| *atxr6$^c$-2* | GTCTGCGAAGAATGCAGTTC |
| | GTTGATGCCACTGATGAACC |
| *agl19$^c$* | GGATAGAGAACGCAACAAGC |
| | GCTAGAGAACTCATAGAGTT |
| *osatxr5$^c$* | GCATGACGTCGGAGATCGAG |
| | GATTTCTTCCGGATTCAGAA |
| *osatxr6$^c$* | GCACTCCAAGAAATCACACG |
| | GCTTGGTCTCCAAGAAGAAG |

**Supplemental Table 6. Primers used in this study**

| Experiment | Sequence (5' to 3') |
|---|---|
| RT-qPCR | |
| *PRX32* | GCTGTAAATTTGGCAGGAGGTC |
| | GCCTTAAGTTGTGGGAGAGTG |
| *PRX38* | ATCTATCGTTTTGGCGGGAGG |
| | TTGTTTAAGTGTAGAAGATGGAC |
| *PRX49* | ACTCCTCTGTTCTTACCGGTGG |
| | AGTGATATCAAGTCCTTGACGG |
| *PRX54* | AGGAAGAAGAGATGGTCTCACC |
| | ACGTATGCGCTCCAGACAAGG |
| *PRX71* | ACACAGTCATTCTCACTCAAGG |
| | CGGAGAATTTCTGTTGTTGAACG |
| *AGL19* | CAGTTAGAGAATCAGTTGGACCG |
| | TCCCATTCCAAGCCACTTC |
| *ATXR6* | CAGACCGATCCTCGTTTCAGTTC |
| | CTTCTTTGACATCACCAAGCTAGTCT |
| *TUB2* | ACTGTCTCCAAGGGTTCCAGG |
| | AAGAACCATGCACTCATCAGC |
| *OsATXR6* | CGCCGTCTTCAGCAACGAG |
| | CGCCATCATCCTCTTGCACAG |
| *OsUBQ5* | AACCAGCTGAGGCCCAAGA |
| | ACGATTGATTTAACCAGTCCATGA |
| ChIP-qPCR | |
| *PRX32_I* | CTACCACAAACTCATTTGCTGCG |
| | ATGTCACGTACTATAATAAATCTATC |
| *PRX32_II* | AGCAACACTGGTTTACCCGACC |
| | TCAACTAAGACGGTCTGGTTACC |
| *PRX38_I* | TGTTTGTTACAATTAAGTTAGGCAAG |
| | CACTATCATGAACACACACAACA |
| *PRX38_II* | CCAATGTCAGTTCATAATGGATCG |
| | CATTGTTTTCTTAGTGTGGCTAGG |
| *PRX49* | GTGGACCAAGTTGGGTTGTTC |
| | AATGGTCTGGAAAGTGTTGTTTGG |
| *PRX54* | TTGACGACACTTCAAGCATCCAG |
| | TTCTCGAGGGCTGTCTTGATAC |
| *PRX71* | TACCAACGGGACGTAGAGATG |
| | CGGAGAATTTCTGTTGTTGAACG |
